# Poisson poisoning as the mechanism of action of the microtubule-targeting agent colchicine

**DOI:** 10.1101/2020.03.25.007757

**Authors:** M. Hemmat, M. Braman, D. Escalante, D.J. Odde

## Abstract

Microtubule-directed anti-cancer drugs, such as paclitaxel, vinblastine, and colchicine, disrupt cell mitosis through suppression of microtubule dynamics (“kinetic stabilization”). However, while the molecular mechanisms of paclitaxel and vinblastine act as pseudo- and true-kinetic stabilizers, respectively, the molecular mechanism of colchicine has remained enigmatic since it requires explanation of both the slow kinetics of the drug and suppression of microtubule dynamics. In this work, we applied an integrated multi-scale modeling-experimental approach to systematically characterize the microtubule targeting agent (MTA) colchicine. We found that colchicine stabilizes microtubule dynamics significantly both *in vivo* and *in vitro* in a time and concentration-dependent manner. Molecular modeling results suggest that tubulin’s binding pocket is accessible to the drug for only 15% of the simulation trajectory time in straight and 82% in curved conformation on average, confirming that colchicine mainly binds to free tubulin. Molecular dynamics simulations show that there are conformational changes at longitudinal and lateral residues of GTP-tubulin-colchicine compared to a lattice tubulin structure, explaining why further incorporation of tubulin dimers to a tubulin-colchicine complex at a protofilament tip is unfavorable. Thermokinetic modeling of microtubule assembly shows that colchicine bound at fractions as low as ∼0.008 to free tubulin can poison the ends of protofilaments with a Poisson distribution and thus, reduce the microtubule growth rate, while stabilizing the tubulin lateral bond and reducing the microtubule shortening rate, i.e. true kinetic stabilization. This study suggests new strategies for colchicine administration to improve the therapeutic window in the treatment of cancer and inflammatory diseases.

**Significance Statement:** Colchicine is an ancient microtubule targeting agent (MTA) known to attenuate microtubule (MT) dynamics but its cancer treatment efficacy is often limited by lack of a detailed understanding of the drug’s mechanism of action. The primary goal of this study was to perform a multi-scale systematic analysis of molecular mechanism of action of colchicine. The analysis indicates that unlike paclitaxel and vinblastine, colchicine poisons the ends of protofilaments of MTs at low fractions bound to tubulin, in a time-dependent manner. Our results suggest new insights into improvement of the clinical administration of colchicine and new colchicine-site inhibitors.

## Introduction

Colchicine, extracted from Colchicum autumnale (1), is a drug first discovered to relieve symptoms of gout based on experimental observations in 15^th^ century (2). In the early 1930s, several studies showed that colchicine had a unique effect on mitosis, arresting cell division at metaphase in tissues both *in vivo* and *ex vivo*, without knowing its primary binding target in cells (3–7). Colchicine studies then led to the discovery of tubulin as the main colchicine-binding protein in mammalian brain, sea urchin eggs, sperm tails, and brain tissue (8–10). It was then that the antimitotic effects of this drug were discovered to be a direct result of its binding to the soluble form of tubulin -subunit of microtubules (MTs)-rather than intact MTs (9–17). As an ancient anti-mitotic microtubule-targeting agent (MTA), colchicine has been the subject of many studies (7, 10, 18–27) to investigate its mechanism of action and assess its potential in cancer treatment, in addition to inflammatory diseases. Colchicine was identified as a narrow therapeutic index drug with unsuccessful cancer treatment clinical trials (28, 29), due to severe toxicity in patients, mainly gastrointestinal problems and multiple organ failures in high does effective for treatment (30). It was suggested that the drug’s high toxicity comes from the slow kinetics of the drug with tubulin with an elimination half-life of 20-40 hours in cells (31). Thus, lower dose colchicine with limited toxicity is an ideal agent for cancer treatment if its efficacy is improved. A recent study investigating gastric cancer both *in vitro* and *in vivo* showed that clinically acceptable doses of colchicine can be promising in inhibiting cell migration, proliferation and tumor growth in nude mice (18, 19). As an alternative, colchicine analogues and other colchicine-site-inhibitors (CSIs) with faster kinetics were taken into consideration for cancer clinical trials (32–34).

At the molecular level, observation of complete mitosis blockage of living cells with only 3-5% of tubulin complexed with 0.1 μM colchicine (7) and MT polymerization inhibition *in vitro* at 1% colchicine to tubulin ratio (7, 20) led to the suggestion of substoichiometric poisoning as the drug mechanism of action. Several studies agree that tubulin-colchicine (TC) complex binds to the tip of a protofilament (PF) of a growing MT and poisons the end of that MT by completely blocking polymerization at the poisoned tip, i.e. an end-poisoning mechanism (21). Alternatively, other studies suggested that the poisoning mechanism is more complicated than a simple blockage of assembly. The alternative hypothesis is that a TC-complex, once incorporated, has a lower affinity for further association of tubulin dimers and that copolymerization of TC-complex with soluble form of tubulin or other TC complexes can occur at the poisoned tip (22, 23). According to this latter mechanism, the poisoning effect (reduced affinity for tubulin) is proportionate to the fraction of TC complexes in the copolymers and occurs at both plus and minus ends (23, 35). Thus, at low TC: tubulin fractions, temporary blockage of polymerization can occur and be repaired by further addition of tubulin and at high TC: tubulin ratios, complete blockage of assembly is observed (22, 23).

Therefore, there is disagreement regarding the lowest fraction required to create complete MT assembly inhibition, mainly due to different experimental methodologies and tubulin source used in previous studies (17, 22, 26, 36, 37). Since incorporated TC-complex caused a reduction in both MT growth and shortening rates *in vitro* (17, 36, 37), it was hypothesized that TC-complex has either a conformational change causing steric hindrance or altered flexibility such that further incoming tubulin dimers are unable to bind to or have lower affinity for a poisoned PF. In addition, the mechanism of the reduction in shortening rate is not well understood (17), but has been proposed to arise from various stabilization scenarios such as strengthening the lateral bonds between PFs (38), altered hydrolysis rate (or Pi-release rate) of guanosine triphosphate (GTP)-tubulin to guanosine diphosphate (GDP)-tubulin (39, 40), or the conformational change induced by colchicine that mimics the more stable conformation of GTP-tubulin (37).

Among the broad range of studies which investigated kinetic rates of colchicine association and dissociation from tubulin (26, 35, 41–44), Garland (42) was one of the first to suggest that colchicine binds to tubulin in a biphasic manner under pseudo-first order conditions. Based on their study, colchicine first binds to tubulin in a relatively fast step (K_1_=6×10^3^ M^−1^), forming the TC complex, followed by a slower step, (k_2_ = (2-3) ×10^3^ s^−1^, k_-2_ = (5-9) × 10^−6^ s^−1^), although the origin of the slow step was not fully investigated. In another study, Banerjee et al. (43) demonstrated that the presence of mixed β-tubulin isotypes with different affinities for colchicine in MTs is responsible for creating the biphasic kinetics, and if each isotype is purified before incubation with the drug, monophasic binding kinetics is observed (43, 45, 46). Despite the disagreements regarding the nature of the colchicine reaction with tubulin, all the previous studies indicate a great deal of consensus over slow binding of colchicine to tubulin as the rate-limiting step in a concentration-dependent manner (7, 10, 16, 21, 44, 47, 48). In addition, colchicine’s penetration into cells has been suggested to be slow (47, 49, 50) (∼1.5-5 as the ratio of internal to external colchicine (7, 51, 52)) although the membrane active agents limiting the drug uptake is mainly unidentified. Colchicine unbinding from tubulin is also established as a very slow reaction, with half-time of dissociation of 10 to 150 hours, for various types of tubulin and experimental conditions (7, 41, 44, 53, 54). The dissociation of TC complex from a MT is also found to be slower than the tubulin itself (53).

Thus, previous studies on the molecular mechanistic details of how the drug works indicate that colchicine characteristics are not well understood at all length-time scales and there are several contradictions that need to be resolved. First, the molecular mechanism underlying the poisoning effect, i.e. reduction in both tubulin association to a TC-capped PF and dissociation of an incorporated TC complex, is yet to be fully understood. Second, there is no direct molecular evidence of why colchicine cannot incorporate directly to MTs. Although the binding site of colchicine to tubulin was published in 2004 (55), little subsequent research has been done to computationally model the TC complex to address the question of how it inhibits MT assembly. Thus, it would be of interest to perform a systematic study of the effects of the MTA colchicine, starting from tubulin at atomistic-level to MT dynamics at cellular level, ideally providing clinically meaningful insights to improve the therapeutic impact of the drug. In the present, we conducted such a multiscale analysis and found that colchicine binds mainly to free tubulin, forming a TC complex, by accessing a binding path with a wide entry channel in soluble tubulin, and cannot access the binding pocket in lattice-incorporated tubulin due to the lattice-induced conformational changes. A TC complex then substoichiometrically poisons the end of PFs by the previously described copolymerization mechanism by which TC complexes cooperatively reduce the affinity of the PF for further tubulin addition. In light of our multiscale modeling results, we identified that a TC complex inhibits MT assembly by having an incompetent conformation for tubulin association at the PF tip and stabilizing lateral bonds responsible for its long-lasting poisoning effect once incorporated. Our results resolve previous seemingly contradictory findings and suggest a cohesive mechanism for suppression of MT assembly by colchicine.

## Materials and Methods

### Thermokinetic modeling

Pseudo-mechanical thermokinetic modeling with added on-rate penalties was used to simulate MT assembly, as previously described (56, 57), using MATLAB R2018a. *In vitro* parameter set (Table S1 in the supporting material) (57) was used to simulate MT dynamics similar to *in vitro* experiments. Colchicine kinetics was simulated based on the following assumptions:

1. Colchicine binding events were limited to free tubulin only since it has a significantly higher affinity for free tubulin compared to polymerized tubulin (1 to 0.001) (11, 14, 17). Thus, a drug binding event in the simulation is defined as binding of a TC-complex to the terminal layer of a PF.
2. All the available colchicine in solution is bound to free tubulin (TC complex) according to our analysis in our thermokinetic model. The analysis indicates that 80% to 100% of colchicine bound to tubulin (TC complexes) would recapitulate the *in vitro* experimental results for 100nM colchicine at 11uM free tubulin concentrations. This result revealed that almost all of the available colchicine in solution is bound to tubulin.
3. A TC-complex will bind to a PF with a probability proportionate to the fraction of TC-complexes to free-tubulin in the solution.
4. A PF is poisoned when a TC-complex is bound to its tip. Poisoning is modeled as a 0.01 fold reduction in k_on_ for the incoming dimers to the PF tip.
5. A 0.05 fold reduction in k_off_ of the TC-complex at the tip is included due to strengthening of the lateral bond when lateral neighbors are present. The lower k_off_ is required to observe the kinetic stabilization of MTs due to a low fraction of PFs poisoned. Otherwise, the TC-complex would unbind before it can stabilize MT dynamics. The alternative mechanism for k_off_ reduction is the strengthening of the longitudinal bond which would result in longitudinal tubulin oligomers in solution. This hypothesis is rejected based on our *in vivo* fluorescent recovery after photo-bleaching (FRAP) experiments in LLC-PK1α cells.
6. The probability of the drug unbinding from a TC-complex is calculated as *p = 1* − exp(−*k*_*d*_ *t*_*min*_), where k_d_ is the drug off-rate (*k*_*off,d*_ *= k*_*on,d*_*K*_*D*_), with K_D_ as the drug dissociation constant, and t_min_ as the current time step from the minimum event time. The k_on_ and k_off_ values were chosen according to previous literature (41–44, 58) and our washout experiments in LLC-PK1α cells.
7. Since the unbinding of the drug from tubulin is slow and it has a low fraction compared to free tubulin, assembly dynamics were simulated for 60 min to observe kinetic stabilization. This is consistent with the *in vitro* experiments were stabilization is observed after a minimum of 30-minute exposure of the drug.

### Molecular dynamics simulation

Molecular dynamics (MD) simulations of free tubulin and tubulin complexed with colchicine were run using NAMD 2.10 software (59) using the CHARMM 36 force field (60) and visualized by VMD 1.9 (61). The protein complex along with the nucleotides were all parametrized using the CHARMM-GUI interface (62). For our study, the tubulin structures were extracted from published tubulin-DARPin room-temperature serial millisecond crystallography (SMC) structures with a high resolution of 2.05 Å (PDB IDs: 5NM5, 5NQT) (63). These structures were chosen over the available tubulin-colchicine-Stathmin-like-domain (SLD) structures (55, 64) because we suspect the existence of the long chain of SLD might cause conformational changes that overshadows colchicine conformations, compared to DARPin-tubulin structure that has lower deviation from an equilibrated tubulin dimer in solution. The simulation system included tubulin heterodimer in two different nucleotide states: GDP, and GTP. Since the structure of GTP-tubulin complexed with colchicine was not available in the same experimental setup, we converted the GDP nucleotide to GTP in the published structures (63) and equilibrated the system. The systems were solvated in TIP3P water using a 10 Å from each side, then neutralized it by adding MgCl_2_ at a 2 mM concentration based on physiologically relevant salt concentrations. Simulations were run in an NPT ensemble (T=310 K, P=1 atm) for a total of 400 to 500 ns for each nucleotide state with and without the drug (1.6 µs). Root-mean-square deviation (RMSD) and root-mean-square fluctuation (RMSF) values were calculated based on tcl scripts written for VMD. BSASA and bending angles were calculated based on the methodology previously described (65).

### Modeling drug transport into the protein binding pocket

In order to identify the transport route that colchicine follows to reach the active site, we used the method, previously described by Escalante *et al.* (2017) (66). In brief, this method uses the motion trajectory of an MD simulation to identify the channels connecting the buried active site of a protein and the solvent exposed surface. The dynamics of the amino acids forming the channel walls are used to identify the most probable residues controlling access into the active site. Finally, we used the same method to determine the likelihood that colchicine is able to reach the active site of tubulin via transport along the identified channels. The last 200ns of the MD simulation trajectories for TC complex and unliganded tubulin in GDP- and GTP-states, along with 200ns trajectory of straight tubulin in solution in both nucleotide states from our previous study (65), were used for this analysis.

### Cell culture

LLC-PK1 porcine epithelial cells stably expressing with enhanced green fluorescent protein (EGFP)–α-tubulin (LLC-PK1α; (67)) or end-binding protein 1 (EB1)-EGFP (EB1/GFP-3; (68)) were cultured in Life Technologies Opti-MEM (Invitrogen, Carlsbad, CA) containing 10% fetal bovine serum (Invitrogen) and frozen in medium containing 10% dimethyl sulfoxide (DMSO) and stored in cryovials in liquid nitrogen before plating. Cells were plated at 50,000 cells/dish in MatTek 35-mm No. 1.5 dishes (MatTek, Ashland, MA), and incubated at 37°C in 5% CO2 overnight before imaging. LLC-PK1α cells were fixed in PHEM buffer (60 mM 1,4-piperazinediethanesulfonic acid [PIPES], 25 mM 4-(2-hydroxyethyl)-1-piperazineethanesulfonic acid, 5 mM ethylene glycol tetraacetic acid [EGTA], and 1 mM MgCl) containing 0.25% glutaraldehyde, 3.7% paraformaldehyde, 3.7% sucrose, and 0.1% Triton X-100, as previously described (69, 70).

### Pharmacological agents

Colchicine (Sigma-Aldrich, St. Louis, MO) was stored as a 100 µM stock solution in deionized water (DI) water at −20°C. PgP inhibitor drug, ly335979 (Zosuquidar), was kindly provided by Dr. William Elmquist (Department of Pharmaceutics, University of Minnesota, MN). Drug stocks was diluted to 2x working concentration in cell culture medium and then heated to 37°C before exchange for half of the cell culture medium. Drug exposure time varied between 30 minutes to 32 hours for colchicine and 30 minutes for ly335979 before the onset of imaging. In control experiments, an equivalent volume of DI water was added to cell culture dishes. Added volume of the drug did not exceed 1% of total solution volume in any condition.

### *In vivo* microtubule dynamics assay

LLC-PK1α and LLC-PK1-EB1 cells were imaged using the Nikon TiE microscope with a 100x/1.49 NA TIRF lens and 1.5× zoom lens (150X total magnification, final pixel sampling size of 65nm), LED illumination (SpectraX; Lumencor Inc., Beaverton, OR), and a Zyla 5.5 sCMOS camera (42 nm spatial sampling; Andor Technology, Belfast, UK). Images for MT dynamics measurements were acquired at 1-s intervals for 1 min with 80% illuminator power and 200 to 300 ms exposure time. Images for estimating hydrolysis rate were acquired by streaming at 500-ms intervals for 20s. During the imaging, the stage and objective were heated to 37°C, with environmental control provided by a BoldLine stagetop incubation system (OkoLab, Pozzuoli, Italy). A PhotoFluor II metal halide light source (89 North, Burlington, VT) and an ET-EGFP filter set (49002; Chroma, Bellows Falls, VT) were used for single-channel GFP imaging. Individual MTs were tracked using the TipTracker semi-automated plus-end localization algorithm without modification (71) with settings enabled for EB1 signal tracking (72). MTs were eligible for tracking if individual plus-ends could be unambiguously tracked (e.g., did not cross another MT) for at least 10 frames (10 s). MTs very close to the edge of the cell were excluded from tracking due to movement confinement. GTP hydrolysis rate was estimated as described previously (72) by fitting an exponential decay function to the EB1-EGFP fluorescence intensity decay in time along the MT lattice. Fluorescence intensity values for fitting were selected from columns (along the time axis) of background-subtracted EB1-EGFP kymographs and normalized to the maximum value. An exponential function was fitted to points starting at the pixel with maximum brightness and terminated at the time point of catastrophe.

Colchicine unbinding experiments were done by first incubating the cells with different concentrations of the drug until saturation effects were observed (30nM after 96 hours, 100nM and 300nM after 24 hours). Then, the drug-containing media was exchanged with Dulbecco’s phosphate-buffered saline (DPBS) buffer for three times and replaced with drug-free media at last. MTs were imaged 8 to 32 hours after washout to search for any dynamic recovery.

### *In vitro* tubulin polymerization assay

Assembly of GTP-Tubulin labeled with Rhodamine (15% labeled, Invitrogen Corp.) onto Rhodamine-labeled biotinylated GMPCPP-Tubulin seeds (7% labeled) was imaged by TIRF microscopy as previously described (73). The Imaging Buffer consisted of BRB80 (80nM PIPES/KOH, 1mM EGTA, and 1mM MgCl_2_) was supplemented with 1 M glucose, 2 mg/ml glucose-oxidase, 2 mg/ml catalase, 1 mg/ml casein. TIRF data were collected using Nikon TiE stand described earlier. Images were acquired at 2 s interval for 10 min. MT end position and backbone coordinates were estimated using TipTracker software as previously described (70, 71) using MATLAB (R2018a). Fiji kymograph tool was also used to generate maximum intensity plots along a user-specified line of interest, here being along a MT growing from the seeds. The slopes of the MT growth vs. tubulin concentration plots were calculated and compared via ANOVA test using *aoctool* function in the statistical toolbox of MATLAB.

### FRAP experiments and estimating free tubulin diffusion coefficient

A spot of ∼1 µm in diameter in the cytoplasm of LLC-PK1α cells were bleached using 488-nm, 100-mW argon-ion laser (Spectra-Physics, Santa Clara, CA) at the Nikon TiE stand described earlier. The TI-PAU attachment and a 50/50 beam splitter were used to direct the laser beam into the rear port of the microscope. The timing of laser illumination was controlled using a Uniblitz VS35 shutter and VMM-TI shutter driver/timer (Vincent Associates, Rochester, NY) set for 2-s delay and 300 ms exposure. Images were collected at 50 ms intervals under 100% power for a total of 10 s (200 frames). The stage and objective were heated to 37°C for the duration of imaging. Fluorescence recovery was measured from time-lapse videos using a custom MATLAB script based on a previously published method (57). Average fluorescence signal was measured for each frame within the bleach region, and a separate, user-defined region distal to the bleach zone to correct for photobleaching that occurs during imaging. In each experiment, intensity values for both regions were normalized to the first 10 frames, and then a normalized, corrected intensity value (I_BC_) was calculated for all frames using Eq. 1.

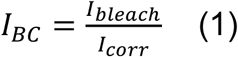

I_bleach_ and I_corr_ are the normalized bleach and correction region intensities, respectively. Next, all frames post-bleach were fit to a single exponential function with two adjustable parameters.

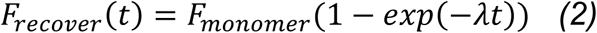

In Eq. 2, F_recover_(t) is the normalized fluorescence intensity at post-bleach time t, *λ* is a constant that determines the timescale of recovery due to the diffusion of free tubulin, and F_monomer_ is the tubulin monomer fraction.

### Statistical analysis

Unless otherwise stated, comparisons between experimental conditions were performed by one-way analysis of variance (ANOVA) or Kruskal-Wallis test with Dunn-Sidak multiple comparisons correction using the anova1 and multcompare functions in MATLAB R2018b and R2019a. Dunn-Sidak multiple comparisons correction was used when comparisons included more than two groups.

### Data and Software Availability

All experimental data, analysis scripts, and simulation codes used in this study are available upon request from the corresponding author.

## Results and discussion

### Colchicine causes kinetic stabilization favoring MT depolymerization *in vivo* in a dose-time-dependent manner

While colchicine is used clinically and has been intensively investigated experimentally, prior studies have not examined the effect of time exposure to the drug as an important parameter in the *in vivo* experiments and have reported ∼100-200 nM as the IC_50_ for most cells (19, 74–77). However, the exposure time of cells to colchicine was ∼24-72 hours in these studies. Therefore, we wanted to examine the dose-time-dependent influence of the drug on single MT assembly using LLC-PK1 cells, up to 32 hours (Fig. 1). To our knowledge, this is the most extensive single MT dynamics analysis for the effects of colchicine over 32 hours. In contrast to paclitaxel and vinblastine where the full effect of the drug kinetics occur in less than 30 minutes of incubation time (57), colchicine binding to MTs was very slow and only longer exposure of the cells to the drug made the MT stabilization effect significant at *nano* molar concentration ranges (up to 8 hours for 30nM colchicine). We used our previously described MT tip tracking and dynamic analysis method (57, 70, 71) to quantify the effect of colchicine on MT assembly dynamics (Fig. 2A). Our analysis in Fig. 2B shows that incubation of colchicine with MTs caused kinetic stabilization, as large displacements of MT tips in both positive and negative directions became less frequent, i.e. the dynamic range of motion of MTs became smaller, with both increasing drug concentration and increasing time of drug exposure. At 600nM colchicine, the MT life history became very similar to fixed MTs within 30 minutes, as the dynamics almost completely disappeared.

**Figure 1.**
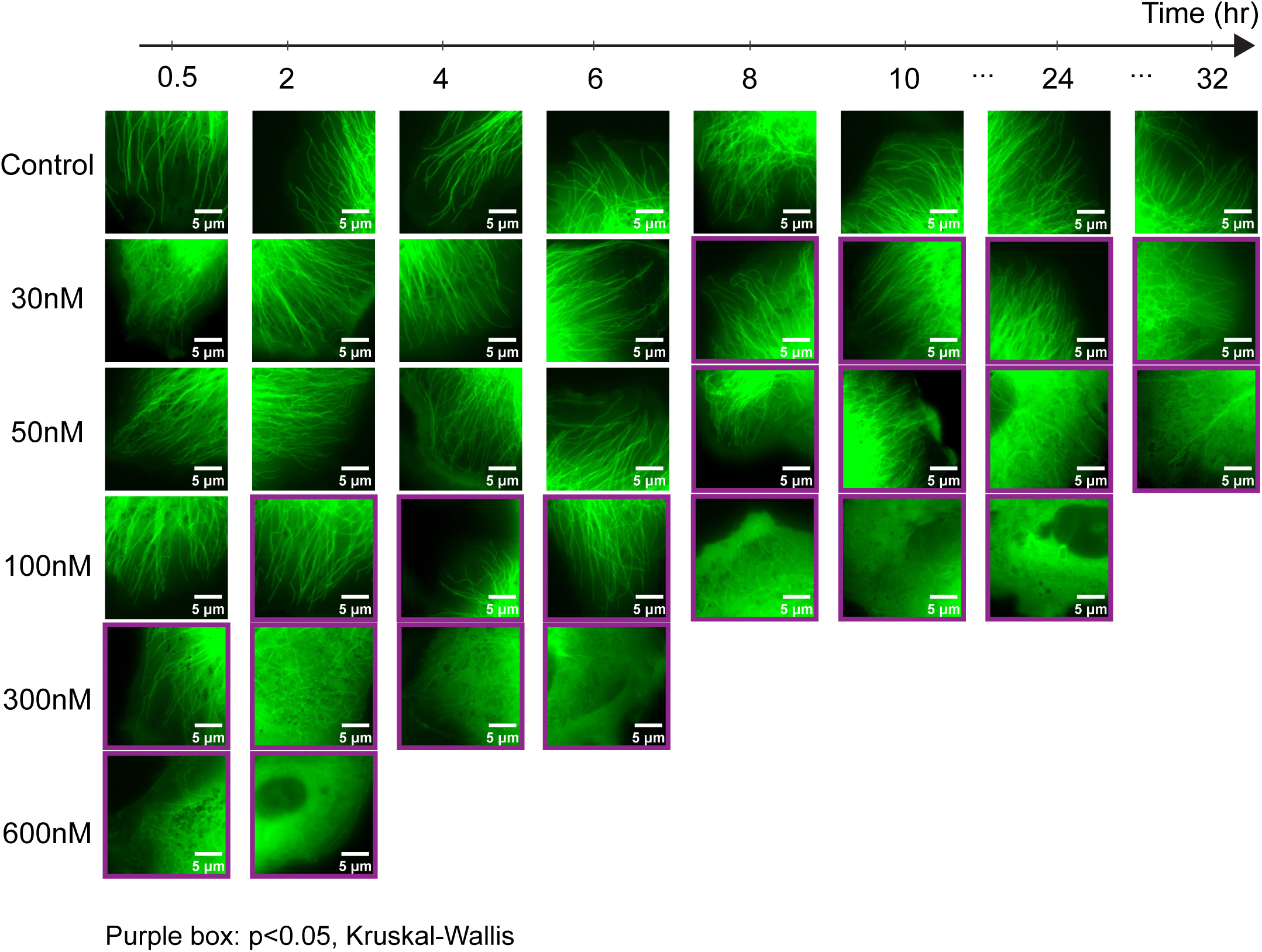
Colchicine suppresses microtubule dynamics in LLC-PK1α cells and promotes MT depolymerization in a time-dose-dependent manner. Individual microtubules are shown in LLC-PK1α cells for control and 30, 50, 100, 300 and 600nM colchicine treated conditions for different incubation time with the drug. Drug-treated cells were imaged until saturating effects were observed with very few MTs. Purple boxes show statistical significance (p < 0.05) in MT dynamics suppression via Kruskal-Wallis test.

**Figure 2.**
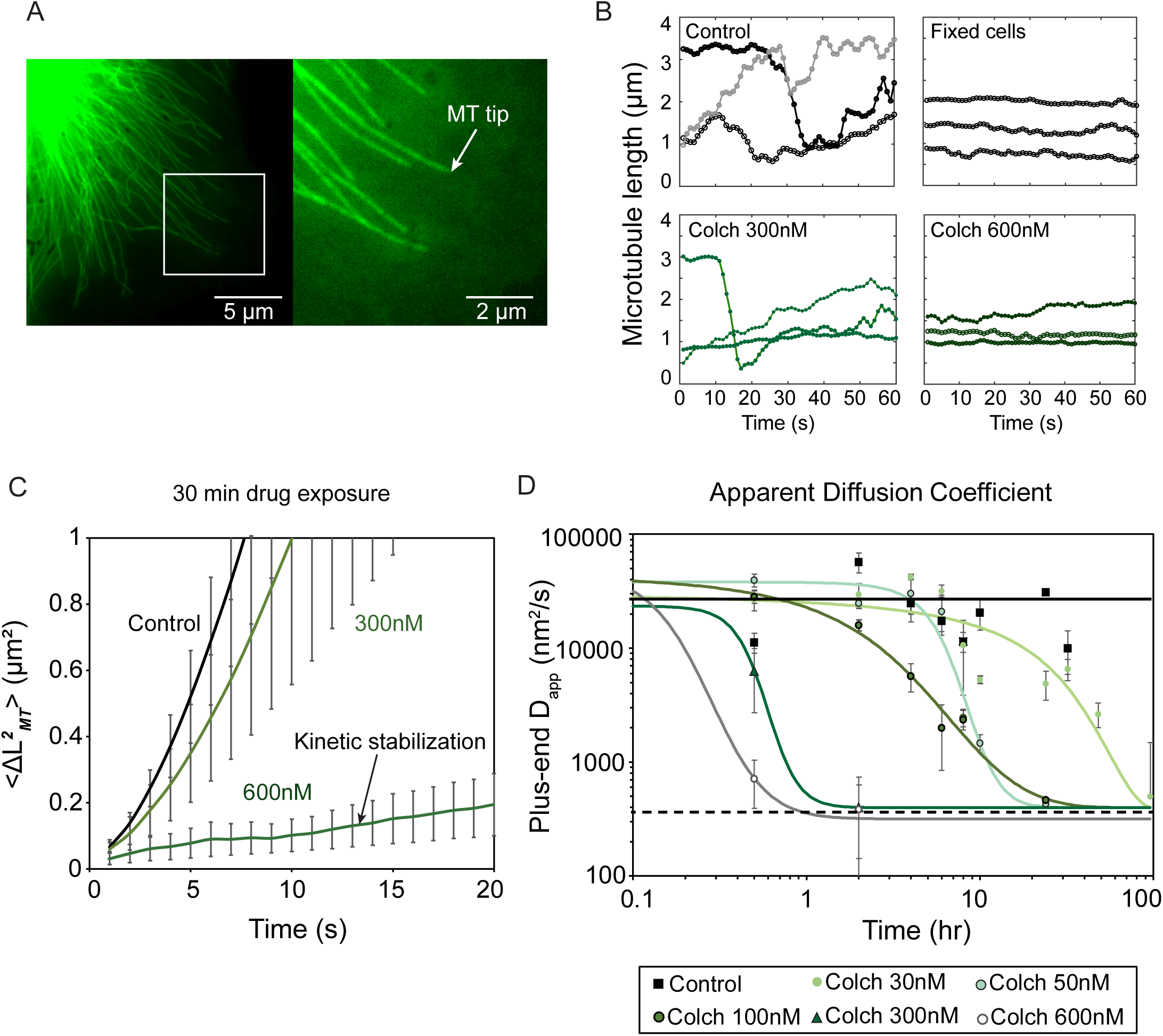
Colchicine reduces MT dynamics in a time-concentration dependent manner in LLC-PK1α cells. (A) An example LLC-PK1α cell is shown with the region (white box) near the cell edge where MTs (white arrowhead) were analyzed. (B) MT plus end position tracked over time for different MTs in control, fixed (see *Materials and methods*), and colchicine-treated cells after 30 minutes exposure. (C) Average MT tip MSD for control cells and cells treated with 300nM and 600nM colchicine for 30 minutes. Plus-end MSD is significantly reduced for microtubules kinetically stabilized by 600nM colchicine due to the loss of dynamic instability. Data points are ± SEM. (D) Apparent MT tip diffusion coefficients (D_app_), estimated from MSD, for various concentrations and exposure time of colchicine. Data points are ± SEM. Curves are best-fit Hill functions. Solid and dashed black lines are estimated diffusion values for live and fixed cells. For each condition, >30 MTs were analyzed from >10 cells.

To examine the kinetic stabilization quantitatively, the diffusion with drift MT dynamics model (78) was used to measure MT plus end apparent diffusion coefficient (D_app_) from the mean squared displacement of MTs (Fig. 2C, 2D). As expected from dynamic behavior of MTs, the diffusion was largest for control MTs (Movie S1 in the supporting material) and reduced with the addition of the drug in a time-concentration dependent manner. Although small MT displacements were still observed at 600nM colchicine, the MT plus end’s diffusion coefficient was significantly reduced even at 30-minute exposure to the drug (Movie S2 in the supporting material). Interestingly, our analysis of MT dynamics for colchicine is different from paclitaxel and vinblastine (57), as the MT tip diffusion with colchicine is strongly time-dependent as well as concentration-dependent while paclitaxel and vinblastine act within minutes at nanomolar concentrations. These results support the notion that time exposure to the drug colchicine is an important part of its binding characteristics besides the dose effects.

### MT dynamics recover ∼32 hours after colchicine washout independent of the colchicine concentration used to suppress dynamics

To determine if the unbinding of colchicine and MT dynamics recovery are also slow similar to binding kinetics, we washed out colchicine from the cells after they had reached the saturation time for kinetic stabilization at various drug concentrations and monitored the recovery of MT dynamics over time. As shown in Fig. 3A and 3B, we found that recovery of MT plus-end dynamics after colchicine washout was similar for all colchicine concentrations used for incubation prior to washout with a recovery time between one to two days. In addition, all the drug-saturated MTs with various colchicine concentrations used prior to washout recovered MT dynamics with a similar rate, i.e. with a half-time of ∼32 hours (Fig. 3C). To assess whether the recovered cells would go through normal migration and proliferation cycles, we imaged them for another 10 to 12 hours post-recovery and found that, similar to control, the cells go through division and migration at approximately normal rates (Movie S3 and S4 in the supporting material). These results confirm that regardless of the initial concentration of the drug, if enough incubation time is allowed with MTs, the drug stabilization effect would be similar at saturation. Our results are also consistent with previous *in vitro* and *in vivo* (7, 37, 41, 44) studies that colchicine binds to MTs with both slow binding and unbinding kinetics and is particularly valuable in understanding high toxicity of the drug due to slow recovery of the binding target.

**Figure 3.**
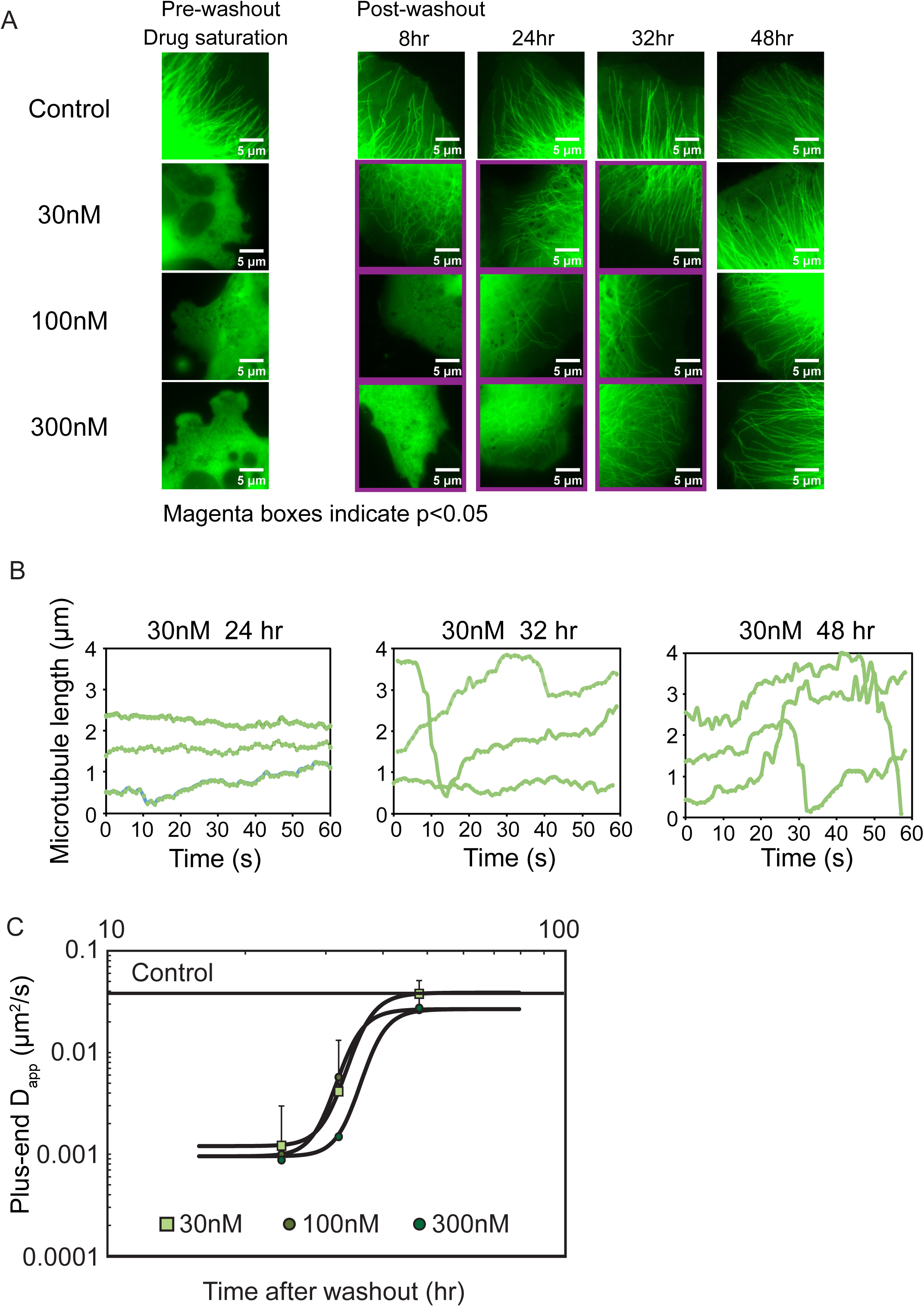
Colchicine washout is followed by slow MT dynamics recovery. (A) Pre-washout control and colchicine-treated cells were washed with PBS and imaged for MT dynamics recovery at 8, 24, 32, and 48 hours after washout. Cells were treated with colchicine (30, 100, 300nM) for various incubation time until saturating drug effects were observed before washout. Magenta boxes show statistical significance (p<0.05) in MT dynamics compared to control cells by Kruskal-Wallis test. (B) MT plus end position tracked over time for 30nM colchicine at different post-washout time points. (C) Apparent MT diffusion is increased with the same rate for all colchicine-treated cells. Solid black line indicates diffusion estimate for live cells.

### GTP hydrolysis rate is not altered by colchicine

Kinetic stabilization by colchicine can be caused by increasing GTP hydrolysis rate which makes it difficult for a MT to maintain a GTP cap required for assembly and converting the MT to a single nucleotide state that is non-dynamic (57). To determine if colchicine suppresses MT dynamics by altering hydrolysis rate, as has been suggested in previous studies (39, 79), we used EB1 as a reporter of the more stable state of tubulin, GTP, and MT growth phase (72, 80). We believe that EB1 comets tagged with EGFP decay following a single exponential function indicating the rate at which GTP-tubulin is hydrolyzed to GDP-tubulin (81). Although some studies show that EB binding to MTs follows a conformational preference in the lattice (82) and might be more complex than recognizing a single nucleotide state, there are evidence showing a strong correlation between MT stability and the size of EB binding region (81). Thus, the comet decay rate at a fixed position on the MT lattice is a marker of the transition rate, k_hyd_, between the stable and unstable states of tubulin in the lattice, as used in several MT assembly models (56, 83–86).

We investigated the EB1 signal decay in control and 300nM colchicine-treated MTs after 24-hour exposure to the drug (saturation). The results in Fig. 4A (Movie S5 and S6 in the supporting material) shows that fewer EB1 comets are observed compared to control and they are more punctate, similar to paclitaxel, vinblastine, and nocodazole (57, 87), due to reduced periods of dynamic growth and shortening. However, the GTP hydrolysis rates, calculated from fitting a single exponential decay for EB1 comets (Fig. 4B), were not significantly altered by the drug colchicine, even though 300 nM colchicine clearly affects MT dynamics (Fig. 4C). This result show that colchicine uses alternative mechanisms besides increasing of the GTP hydrolysis rate to create kinetic stabilization *in vivo*.

**Figure 4.**
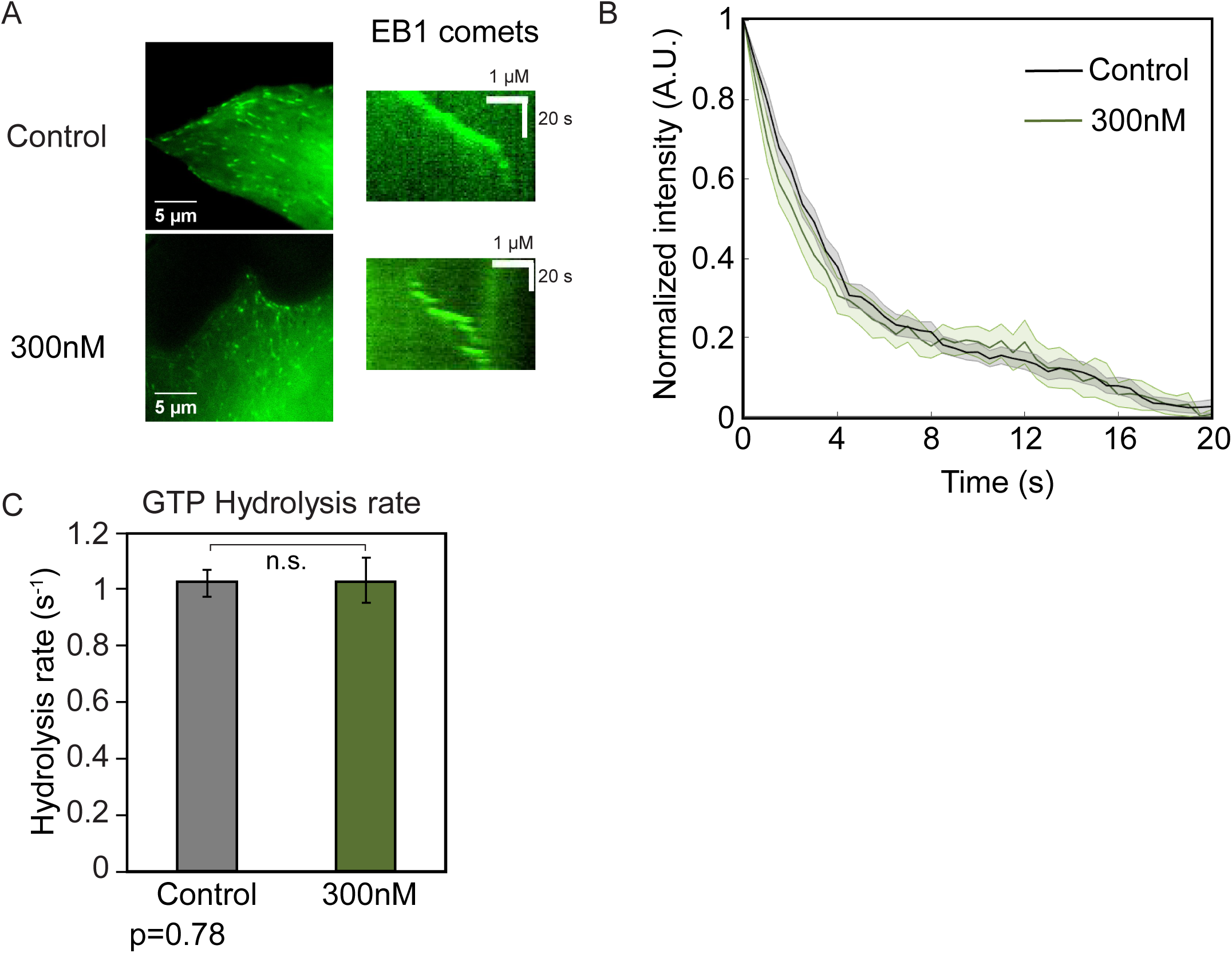
Colchicine does not affect GTP hydrolysis rate *in vivo.* (A) Example LLC-PK1 cell region stably expressing EB1-EGFP (left) and example kymograph for an individual EB1 comet for control and 300nM colchicine treated cells (24-hour exposure). (B) average normalized fluorescence recovery curves for control (black) and colchicine-treated (green) cells. Error bars are ± SE. (C) Hydrolysis rate, as estimated from the best-fit exponential decay rate of EB1-EGFP signal, is not altered by colchicine treatment. Error bars show mean ± SEM. *p<0.05 by Kruskal-Wallis test.

### Theoretical mechanisms for MTAs: True- and pseudo-kinetic stabilization

Two theoretical mechanisms for kinetic stabilization of MTs by MTAs were found by a previous study (57) for the MTAs vinblastine and paclitaxel: true- and pseudo-kinetic stabilization, respectively. In this framework (57), the MTA can significantly (more than 10-fold) affect on- and off-rates (k_^*^on,PF_, k_off,PF_) to attenuate dynamic instability, i.e. MT net assembly rates and plus-end diffusion coefficient, which is characteristic of a true kinetic stabilizer (tKS) such as vinblastine. As the alternative mechanism, the MTA can minimize the ΔΔ*G*^0^, the energetic difference distinguishing between the stable (GTP) and unstable (GDP) states of tubulin, without affecting the kinetic rates of assembly. This mechanism can be achieved through an ensemble of conformational changes in tubulin structure that makes GTP-state more like GDP-state or vice versa. This will result in the suppression of MT dynamics due to having a single state tubulin, which is characterized as a pseudo-kinetic stabilizer (pKS), such as paclitaxel. Since colchicine creates a similar phenotype in MTs, we examined its effects on the assembly dynamics within this framework. We chose this framework because: 1) the mechanisms were defined according to the experimental observations’ constraints and a computational model of MT assembly (56) with five independent parameters describing MT dynamics: 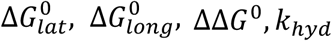 *and k*_*on*_ (Table S1), recapitulating MT dynamics consistent with a range of experimental observations (56, 57, 78), and 2) Each of the parameters of the model can be tested through various *in vivo* and *in vitro* experiments and molecular simulations for effects on dynamic behavior of MTs.

### Colchicine reduces MT growth rates *in vitro*, consistent with a true kinetic stabilization mechanism

In order to determine whether colchicine significantly affects the association and dissociation rates of assembly, we used the *in vitro* MT assembly assay where kinetic rates are estimated based on growth rate (v_g_) measurement as a function of free tubulin concentration ([Tub]) (57, 78). Since the slope of this plot would be proportional to the sum of kinetic rates based on a 2-D model (78), we predict that a tKS would significantly reduce the slope, as in the case of vinblastine, whereas a pKS will not have a significant effect on the rates, as in the case of paclitaxel. Colchicine at high concentrations (100nM) stabilized MTs with some nanoscale dynamics observed after 30 minutes exposure, and at low concentrations (30nM) did not alter the dynamics significantly compared to control MTs for 11 µM free tubulin concentrations (Fig. 5A-5D). At 50nM concentrations, we observed short periods of assembly inhibition, occasionally recovered during the imaging period (10 minutes) (Fig. 5C). Growth rates, measured using kymographs of the plus-ends of MTs *in vitro* (Fig. 5E), decreased with increasing colchicine concentrations and shortening rates were not affected remarkably (Fig. 5F), in line with previous *in vitro* studies (37, 88). We found that colchicine reduces the slope of growth rate vs. [Tub] by ∼3 fold (∼10 to 20 fold change in the kinetic rates (57)), thus, making it a tKS, similar to vinblastine (Fig. 5G). The reduction in the kinetic rates can be either achieved through a change in the lateral or longitudinal bond energies in the model, as described by Castle *et al.*, 2018.

**Figure 5.**
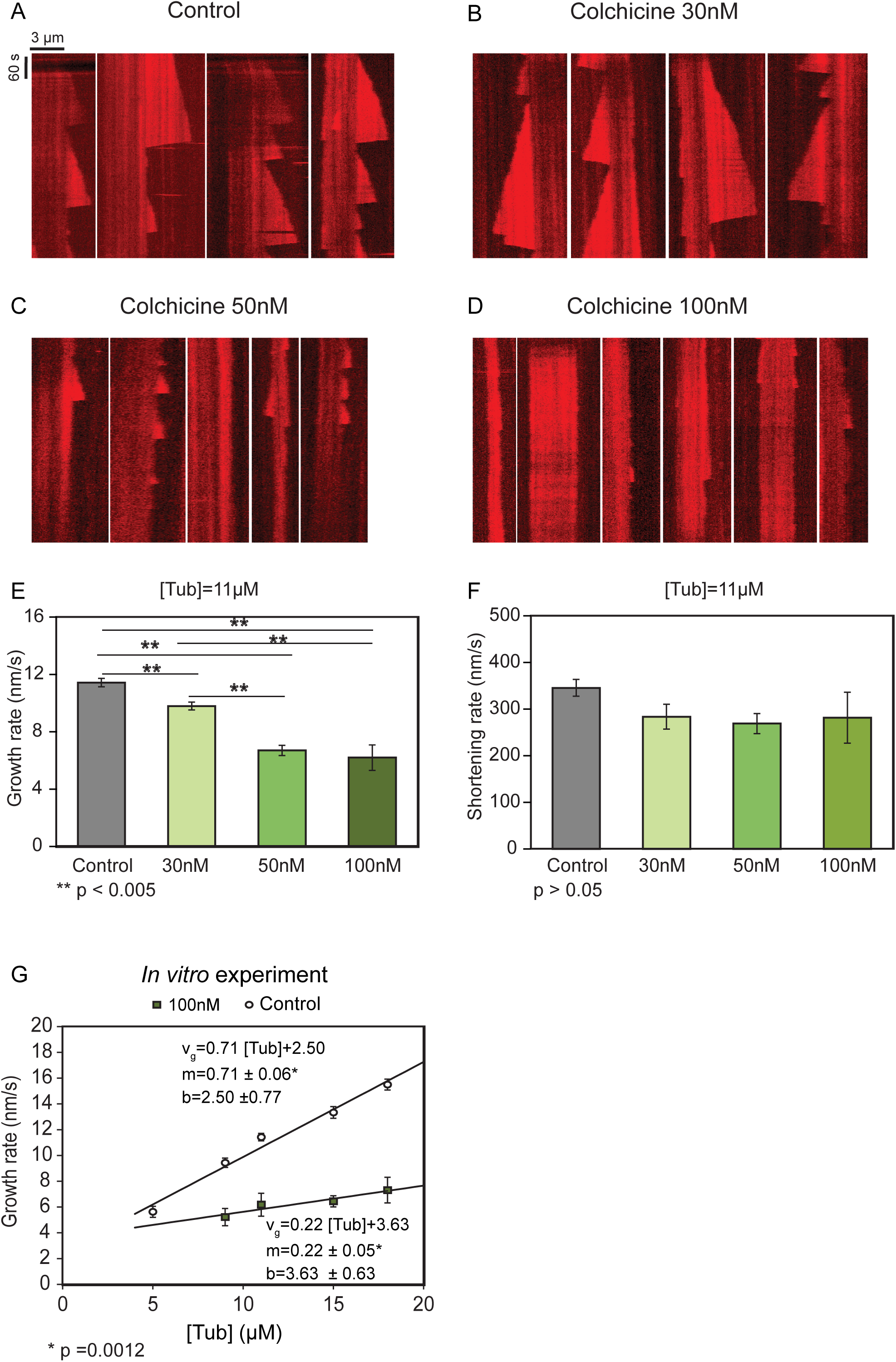
*In vitro* MT growth rates are decreased by colchicine. Example kymographs of MT growth from stable GMPCPP-seeds *in vitro* for (A) control, (B) 30nM, (C) 50nM, (D) 100nM colchicine. *In vitro* experimental estimates of average MT (E) growth rates and (F) shortening rates vs. colchicine concentration. Error bars are mean ± SEM. **p<0.005 compared to control by Kruskal-Wallis, corrected for multiple comparison. (G) MT growth rates for a range of free tubulin concentrations shows significant decrease in slope for 100nM colchicine, consistent with a tKS mechanism. Slope (m) and intercept (b) are estimated from linear best fit ± SE. *p=0.0012 compared to control by co-variance analysis.

Note that although we did not observe any effects on MT dynamics *in vivo* at 100nM colchicine after 30 minutes exposure, we detected significant stabilization effects on dynamics at 100nM *in vitro*. This is likely because of reduced permeability (influx) or efflux of the drug out of the cell membrane *in vivo* which requires higher concentrations of the drug to see similar effects to *in vitro* (49, 89, 90). However, higher concentrations of colchicine would mean stronger stabilization effects, and even at 100nM, the slope is significantly reduced *in vitro*. Therefore, our conclusions of colchicine being a tKS based on our *in vitro* experiments remain valid. We also note that colchicine binding to tubulin is a slow process (37, 48), thus, not all the available colchicine might have formed TC complexes after 30 minutes of incubation with tubulin and the observed effects might have been under or close to saturation.

### PgP-inhibition does not affect the uptake of colchicine in cells

To further investigate the effective colchicine concentration inconsistency between *in vitro* and *in vivo* experiments, we tested the hypothesis that efflux pumps in our cells are excreting part of the permeated drug out of the cell membrane. We inhibited the P-glycoprotein (PgP) pump due to being a strong substrate for colchicine among all efflux pumps (90–92) by a potent PgG inhibitor, LY335979 (Zosuquidar) (93, 94) in our cells. The cells were then incubated with the drug colchicine and examined for effects on MT dynamics (Fig. 6A, 6B). The results first confirmed that LY335979 at 1µM did not alter normal MT behavior, by itself. Furthermore, colchicine at concentrations with saturating effects *in vitro* (50 and 100nM) did not have any significant effects on plus-end diffusion of MTs *in vivo* after PgP inhibition (Fig. 6C). Our results are consistent with the finding that in LLC-PK1 wild type cells PgP expression is low (95), and even if the expression is altered in our α-tubulin expressing cells (LLC-PK1α), it is not involved in excreting colchicine. The alternative hypothesis for lower colchicine efficacy in our cells is that the permeability in the membrane is the limiting factor in the entrance of the drug which reduces the effective intracellular concentration of colchicine *in vivo*.

**Figure 6.**
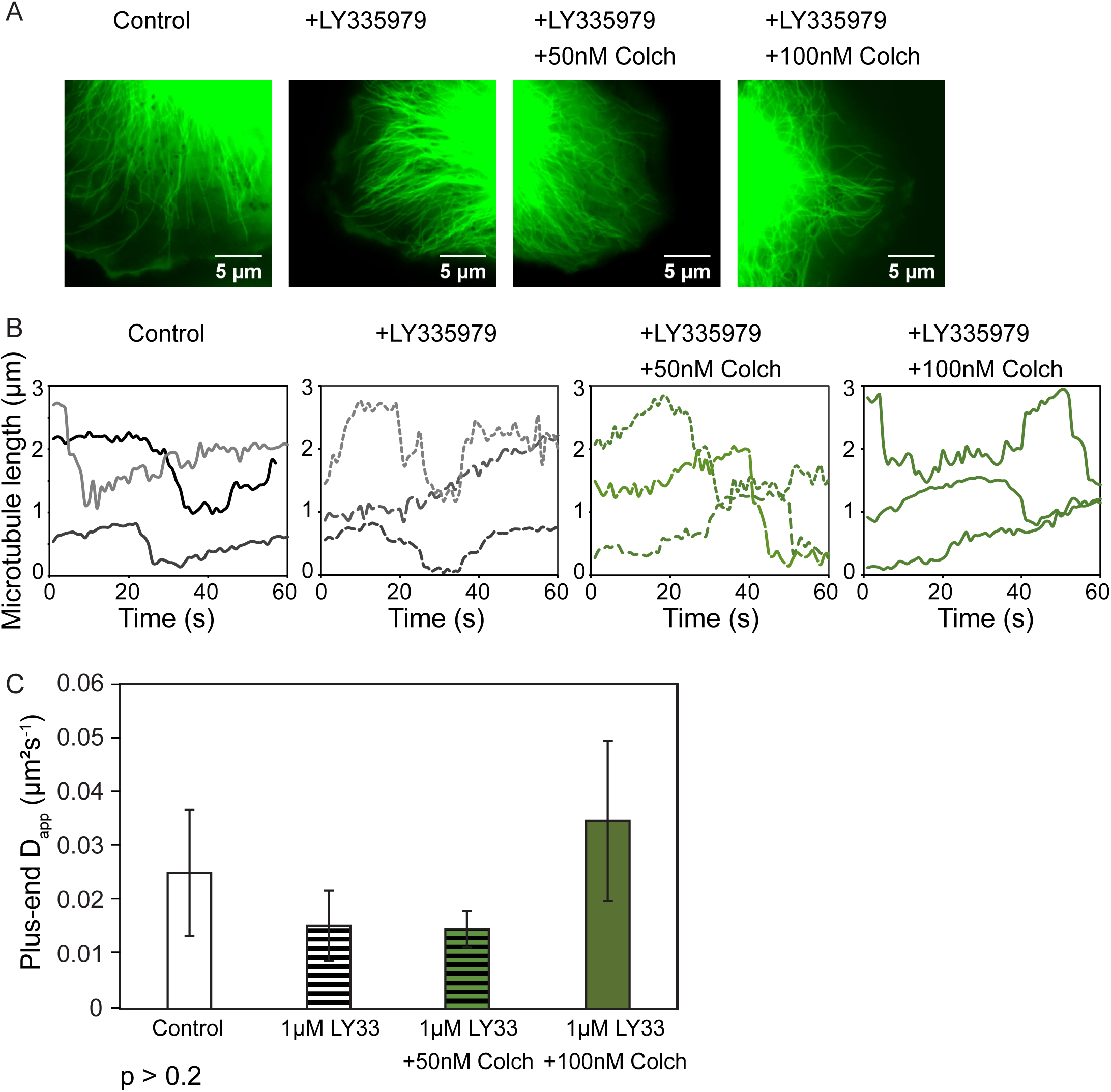
PgP inhibition does not affect colchicine stabilization effects in LLC-PK1α cells. (A) Example analysis region of LLC-PK1α cells are shown for control cells and cells treated only with PgP inhibitor, Ly335979, or treated with LY335979 and colchicine afterwards (50nM, and 100nM). (B) MT length-time histories tracked over time for different drug-treated cells after 30-minute exposure. (C) MT apparent diffusion coefficient is not modified by addition of Ly335979 or colchicine concentrations of 50 and 100nM. Error bars are mean ± SEM. p>0.2 by Kruskal-Wallis test, corrected for multiple comparisons.

### Unlike vinblastine, colchicine does not promote tubulin longitudinal oligomers in solution

Based on our *in vitro* results, colchicine slows the kinetic rates of tubulin assembly into MTs through changing lateral and/or longitudinal bond energies. To detect whether the longitudinal bond is stabilized, similar to vinblastine, we determined if longitudinal oligomers are found more frequently in the cytoplasm with addition of colchicine. The presence of the oligomers is found by measuring the diffusion coefficient of free tubulins in the cytoplasm, as higher molecular weight species have lower diffusion rates (57). Thus, we bleached a region in the cytoplasm of LLC-PK1α cells treated with saturating concentration of colchicine after 24 hours of exposure (Fig. 7A) and monitored the EGFP signal recovery (Fig. 7B). The diffusion coefficients calculated based on FRAP equation or exponential fit (see Material and methods) indicated that colchicine did not affect the average diffusion coefficient of free tubulins (Fig. 7C). Hence, in contrast to vinblastine (57), colchicine does not promote oligomerization, consistent with previous studies (64, 96), and did not strengthen the longitudinal bond. Note that lateral oligomers cannot be detected in the solution due to the lateral bond being weak with a short lifetime (65, 97). These results altogether provide evidence that colchicine stabilizes dynamics by strengthening the lateral bond in the MT thereby slowing off-rates of tubulin bound to colchicine in the MT lattice. However, further investigation is required to explain the reduction in the on-rate constant predicted for TC complex because lateral bond stabilization by itself cannot induce slow binding kinetics.

**Figure 7.**
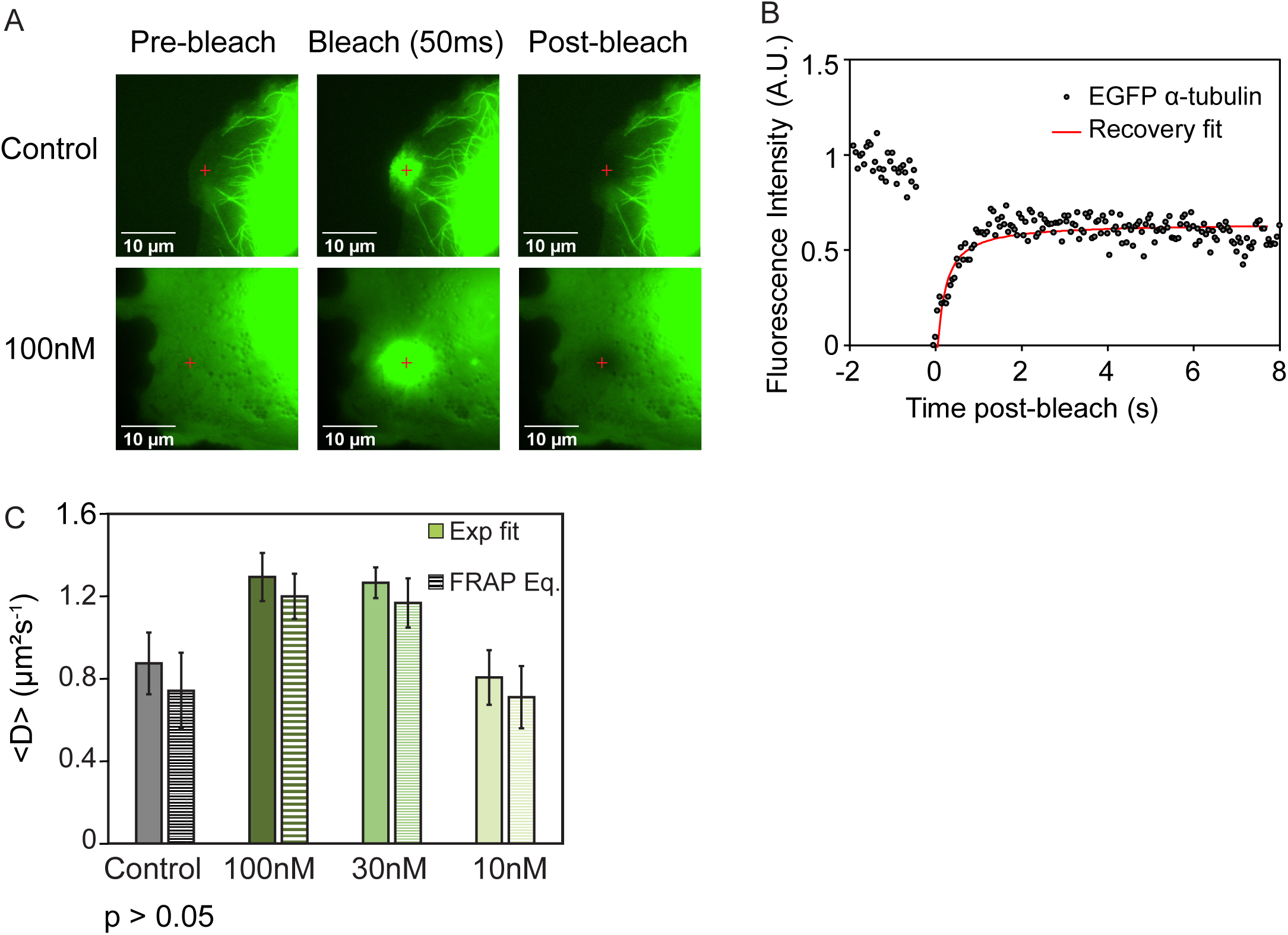
Colchicine does not promote tubulin oligomerization in LLC-PK1α cytoplasm. (A) Example cytoplasm region at the periphery of the cell is shown before and after photobleaching for control and 100nM colchicine treated cells. (B) Normalized EGFP-α-tubulin fluorescence signal is shown pre- and post-bleaching in a cell. Red line indicates best-fit exponential curve to the recovery data. (C) Average diffusion coefficients of tubulin for control and drug-treated cells were not significantly different. p>0.05, obtained by Kruskal-Wallis test, corrected for multiple comparisons.

### Molecular dynamics simulations reveal conformational changes at lateral and longitudinal residues of GTP-tubulin-colchicine complex

Thus far, we have shown that TC complexes poison the PF ends that they bind to by reducing the association rate of further tubulin dimers at the PF tip. To investigate if there is a conformational change, bending flexibility or bending preference change reflected at atomistic and molecular level, we performed MD simulations of free tubulin dimers unbound and bound with colchicine with GDP or GTP nucleotides (Fig. 8A). The MD trajectories were analyzed for any significant change in bending angles, dimer flexibility, dimer compaction, and intradimer interactions of colchicine-bound tubulin (TC) compared to free tubulin (Tub), based on the mechanisms suggested by previous studies (25, 42, 58). GDP-colchicine bound structures demonstrated steady state RMSD of their backbone atoms after ∼100ns (Fig. 8B), while GTP-state showed higher RMSD on average compared to GDP-state and required longer simulation time to reach steady state. Note that based on the hypothesis that colchicine can only incorporate into the MT lattice via TC complex formed with a GTP-nucleotide, and not via addition of colchicine directly to lattice-bound tubulins, we were mainly interested in the conformational changes in the GTP-state TC structure.

**Figure 8.**
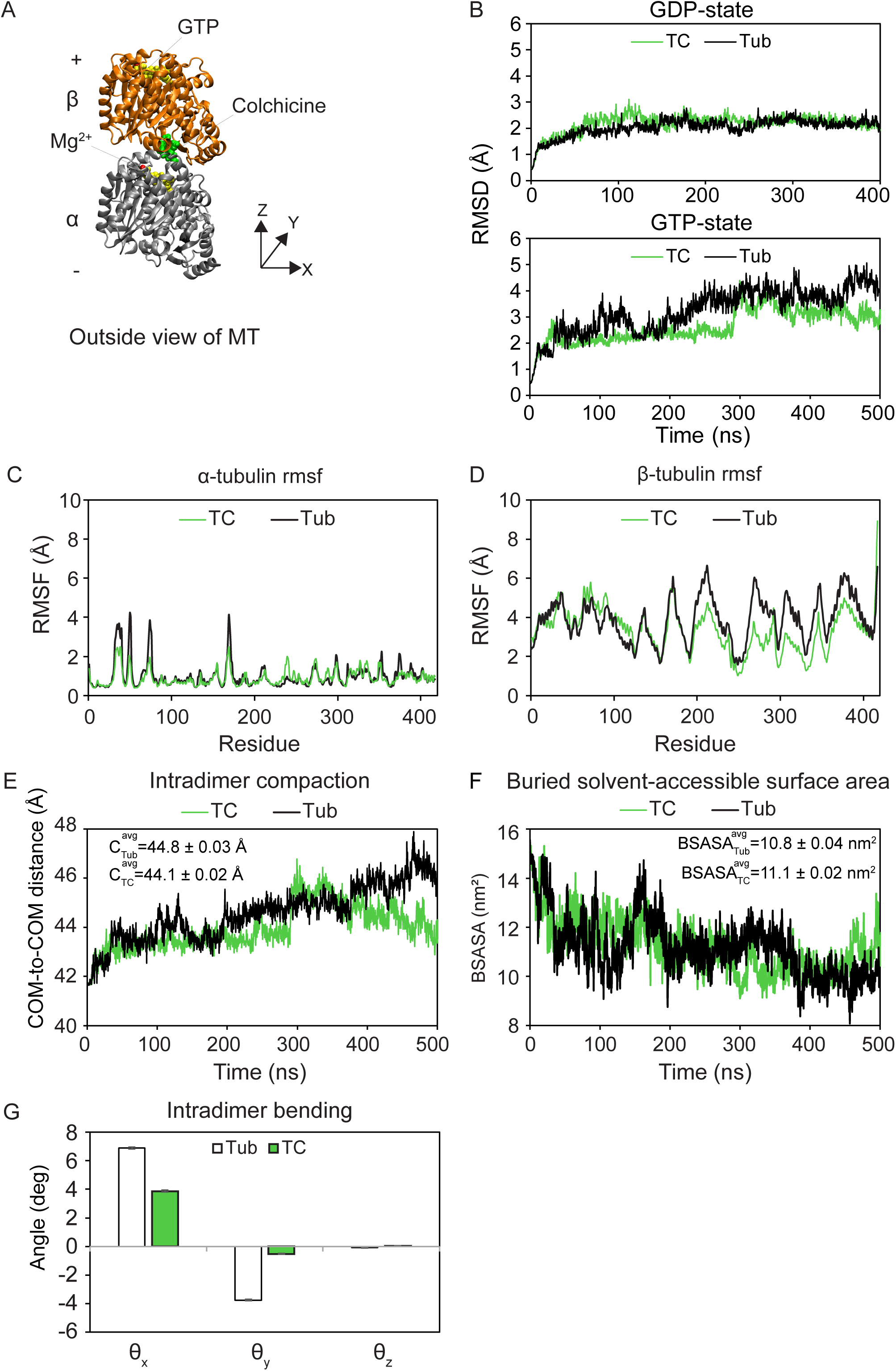
Molecular dynamics simulations revealed structural changes caused by colchicine binding in tubulin. (A) Simulation system is composed of one tubulin heterodimer bound and unbound with colchicine in GDP- and GTP-state. (B) RMSD of backbone atoms over time is depicted for TC and Tub complexes in GDP- (top) and GTP-states (bottom) for 400ns and 500ns trajectories, respectively. Average RMSF values of (C) α- and (D) β-tubulin for GTP TC and Tub complexes shows lower flexibility for colchicine bound tubulin. (E) Intradimer compaction, estimated from COM-to-COM distance of α- and β-subunits, is higher for GTP-TC complex compared to Tub. (F) Hydrophobic interactions, estimated from BSASA, are stronger in GTP-TC complex compared to Tub due to a more confined flexibility and motion. (G) Intradimer bending angles are lower on average for TC complex compared to Tub due to limited flexibility in motion of the residues. Error bars are average of the boot-strapped data ± SEM.

Colchicine is hypothesized to alter tubulin dimer flexibility, thus, making the dimer unfavorable for fitting into a straight MT lattice and blocking the polymerization consequently (25). Our MD results provide evidence in favor of a decreased flexibility in GTP-TC complex compared to the unliganded tubulin (Fig. 8C and 8D). By contrast, binding of colchicine to GDP-tubulin did not significantly change the average fluctuations of the tubulin residues (RMSF; Fig. S1A and Fig. S1B in the supporting material). The results indicate a nucleotide-dependent conformational change in tubulin as a result of colchicine binding where the flexibility of a GTP-tubulin is reduced upon colchicine binding.

We also considered the possibility that colchicine changes the intradimer compaction of tubulin, making it inconsistent with lattice spacing and thus, any incoming dimer would have perturbed lateral and longitudinal bonds. Surprisingly, both Tub and TC dimers relaxed their initial intradimer compaction within 20 ns of simulation; however, TC dimer in GTP-state maintained a higher intradimer compaction (by 0.7 Å on average) compared to the unliganded tubulin (Fig. 8E). By contrast, GDP-state TC complex showed a consistent lower dimer compaction, ∼0.5 Å on average, than the free dimer (Fig. S1C in the supporting material). As larger intradimer compaction means weaker hydrophobic interactions, buried-solvent-accessible surface area (BSASA) was calculated for both dimers and GTP-TC complex showed stronger hydrophobic interactions (Fig. 8F) while GDP-TC complex indicated smaller BSASA, by 1.7 nm^2^ on average, as expected (Fig. S1D in the supporting material).

A conformational change can also be reflected as a change in bending preference of the TC complex relative to the free tubulin. Based on previous MD studies (65, 98, 99), free tubulin conformation is bent and moves toward a more bent conformation if simulated long enough. To test this hypothesis, we measured the intradimer bending angles of the dimers (Fig. 8G, and S1E), decomposed in radial (θ_x_), tangential (θ_y_) and twist (θ_z_) bending directions (65). Both dimers in solution moved toward a more bent conformation, as predicted by previous MD studies, in both radial and tangential directions with GTP- and GDP-TC complexes being more confined in bending motions compared to the unliganded tubulin dimer.

Finally, to address the question of how divergent the TC complex structure is from an unliganded tubulin structure in solution, we calculated residue-by-residue RMSD of the TC complex compared to Tub structure, both averaged through the last 200ns of the trajectory (Fig. 9A and 9B). Interestingly, GTP-TC complex structure showed higher divergence around the longitudinal interfaces and both lateral interfaces, while GDP-TC complex was very similar to its unliganded structure. These results showed that the major conformational changes occur in the GTP-state tubulin which is the nucleotide state of an incoming TC dimer to a PF tip. In addition, those conformational changes are located around the longitudinal residues, suggesting the possibility that they are responsible for making the TC complex less compatible for further dimer addition. In addition, the observed changes around lateral residues, suggests that they might be responsible for strengthening the lateral bond (slowing the dimer dissociation).

**Figure 9.**
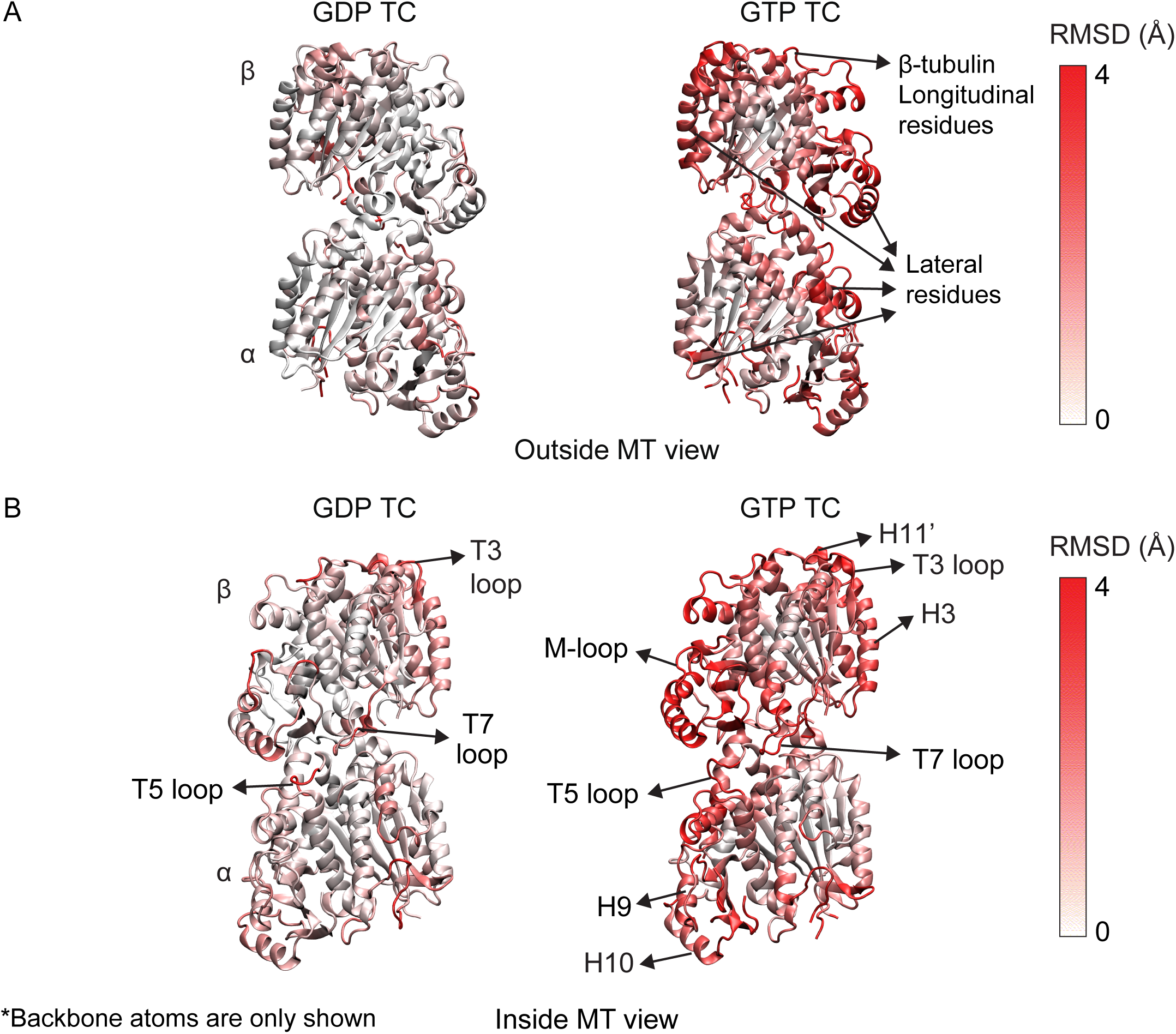
Residue-by-residue comparison of the average TC complex structure compared to unliganded tubulin structure (Tub) in GDP- (left) and GTP-states (right) with (A) MT outside view and (B) inside view. RMSD is calculated from comparing average TC structure to average Tub structure. Longitudinal and lateral bond zones indicate higher deviations from unliganded tubulin in GTP-state. Color bar shows RMSD values in Angstrom. Residues with high RMSD involved in lateral and longitudinal bond are highlighted with arrowheads. Color bar shows RMSD values in Angstrom. Backbone atoms are only shown for better visualization.

### Colchicine can mainly bind to free tubulin in a curved conformation

Previous studies have identified colchicine’s preferential binding to the soluble form of tubulin rather than polymerized tubulin (11, 14), presumably due to the straight tubulin structure in a MT lattice (55, 100). To further explore this idea, we took advantage of the available structures of colchicine-bound tubulin (TC), unliganded soluble tubulin (Tub), and polymerized tubulin (Tub_s_) to quantify the accessibility of the binding channel of colchicine, as previously described (66). We used drug transport modeling method for colchicine transport because it has successfully predicted the transport of contaminants through several degradation enzymes (101), and ligand transport along hydrophobic enzymes (102), with binding channels similar to the hydrophobic binding pocket buried inside tubulin.

As shown in Table 1 for the GTP-state tubulin, the average channel length varies significantly depending on the conformation of tubulin. GDP-state results were similar to GTP-state (Table S2 in the supporting material). Colchicine would have to travel the longest distance in the polymerized form (38Å), whereas in the unpolymerized form, the active site of tubulin is relatively close to the surface of the protein (9.2Å). Similarly, there is a large difference in the average bottleneck radius, where the tightest channel was observed in the Tub_s_, being just 1.3Å. On the other hand, both Tub and TC showed an average bottleneck radius of ∼3.5Å.

**Table 1.** Average length and radius of the identified channels connecting the surface and buried active site of GTP-tubulin. The (R) and (L) refer to the direction of the channel as defined in Fig. 10.

|  | Average<br>Channel length (Å) | Average bottleneck<br>radius (Å) | Colchicine<br>entrance (%) |
| --- | --- | --- | --- |
| Tub (L) | 9.2 ± 0.5 | 3.6 ± 0.5 | 82 |
| TC (L) | 23 ± 2 | 3.1 ± 0.1 | 71 |
| Tub<sub>s</sub> (R) | 38 ± 1 | 1.3 ± 0.1 | 15 |

It is important to note that the channels identified in the polymerized form of tubulin point in a different direction than the ones found in TC and Tub. We have shown these two different channel directions in Fig. 10A and refer to them as R (right) and L (left), for the respective channels found in the polymerized and soluble forms of tubulin. We found that in Tub_s_, the L channel is not capable of being formed due to the movement of the loop containing residues 244 to 260 (T7 loop) of the β-subunit, shown in Fig. 10B, as suggested by Ravelli *et al.* (2014). We were able to identify the L channel in Tub_s_ only in <0.1% of the analyzed MD simulation frames. The movement of the T7 loop in β-subunit results in the formation of the R channel. The R channel was never identified in any of the analyzed frames for TC and Tub structures.

**Figure 10.**
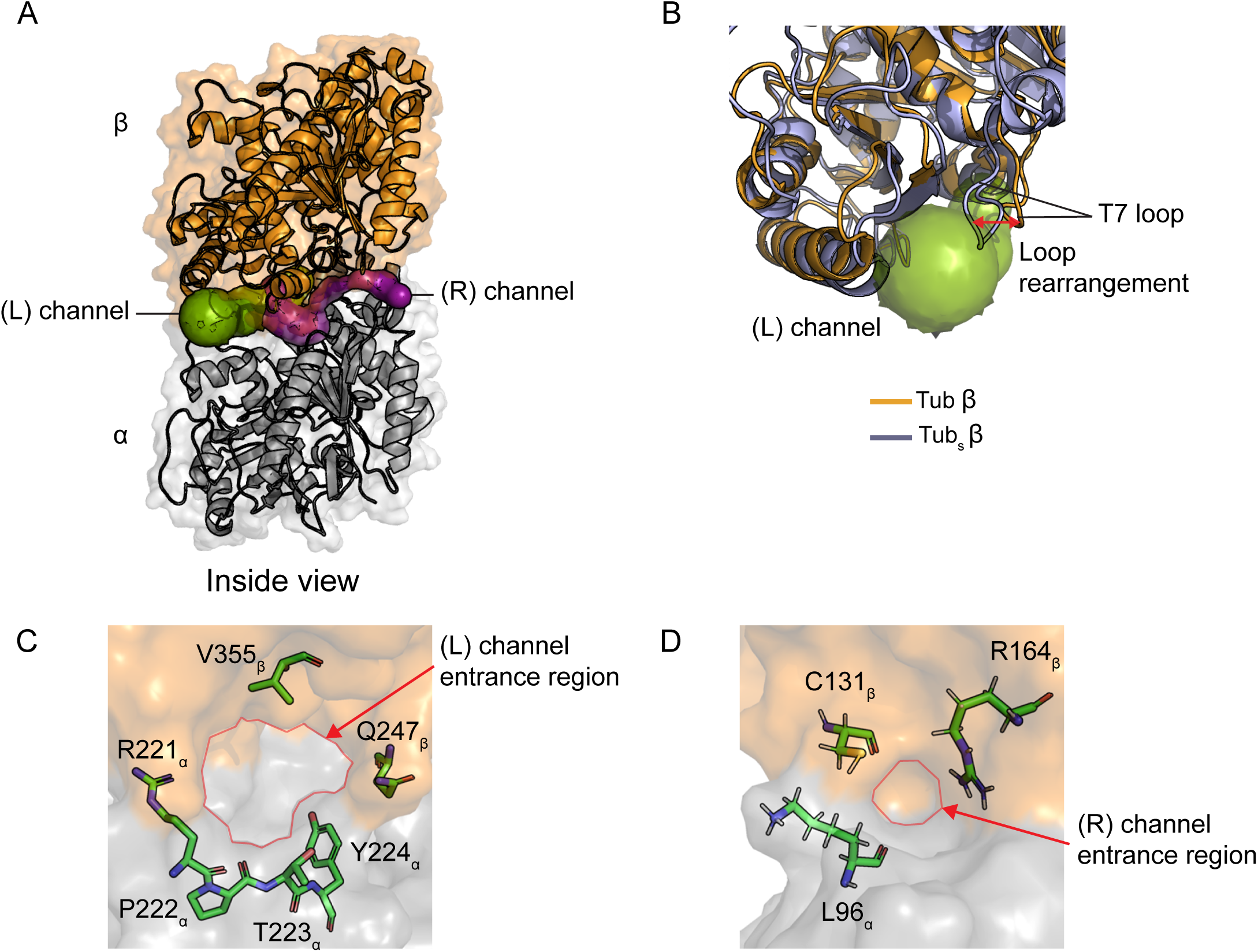
Colchicine binding channel, estimated from connecting the buried active site of tubulin and the solvent exposed surface, is more accessible to the drug in curved tubulin structures (TC and Tub) compared to a lattice straight tubulin structure (Tub_s_). (A) Binding channels identified for colchicine in curved tubulin (L channel) and straight tubulin (R channel). The L and R channels shown were not identified in the same frame, they have been overlaid for illustrative purposes. Inside view of MT is shown. (B) Overlay of the β-subunit loops in the unpolymerized (Tub-orange) and polymerized (Tub_s_-purple) forms of tubulin showing the movement of the β-T7 loop containing residues 244-260. The residue showing the largest movement is Q247 with a maximal displacement of 7.1Å between the two forms. The displacement of these residues causes the L channel to be fully closed in the polymerized from of tubulin. (C) Entrance region of the L channel and (D) R channel in Tub and Tub_s_ structures. The residues shown form the narrowest portion of the channel, i.e. bottleneck. The red line indicates the entrance region for colchicine which is significantly smaller in the Tub_s_.

The third component of our analysis was to calculate the frequency at which colchicine was able to move through the tunnels and reach the active site, i.e. successful binding of colchicine to tubulin. As shown in Table 1, colchicine is most likely to reach the active site in the soluble form of tubulin, Tub, closely followed by TC complex. It is important to note that the transport of colchicine was calculated based on the identified L channel structures. Interestingly, we found that colchicine is only able to reach the active site of Tub_s_ for 15% of the trajectory time. The major hindrance to the movement of colchicine through the channels is imposed by two different sets of residues, one set for each of the L and R channels, respectively, as shown in Fig. 10C and 10D.

The different channel pathways identified in the Tub_s_, TC and Tub structures are the result of the movement of a single loop (β-T7). However, this seemingly minor change in conformation causes a major change in the transport properties of colchicine into the active site of tubulin. These results provide evidence that colchicine is unable to bind to the polymerized form of tubulin due to the binding channel inaccessibility and mainly binds to soluble form of tubulin in a nucleotide-independent manner, as previously suggested by several studies.

### Thermokinetic modeling predicts substoichiometric Poisson poisoning of microtubule assembly by colchicine

We used a thermokinetic model of MT assembly (56), modified with on-rate penalties for lagging PFs (57), to study the possible mechanisms for colchicine-mediated kinetic stabilization constrained with experimental observations. Our molecular modeling confirmed previous reports that colchicine preferentially binds to free tubulin and has a weak affinity for MTs (11, 14, 17). Thus, we only allowed drug addition to free tubulin, i.e. forming TC complexes. We then simulated the on- and off-kinetics of TC complexes at the tip of a MT (see Material and methods). The poisoning effect of colchicine as a tKS was implemented as a reduction of both association onto and dissociation of the TC-complex from the PF end (Fig. 11A). We believe that an absolute blockage of polymerization (simple end-capping mechanism) by colchicine is not consistent with our *in vitro* experimental results, indicating occasional repair and recovery of dynamics at lower concentrations (50nM, Fig. 5C). To determine the drug kinetic parameters that best reproduce the stabilizing effects on dynamics *in vitro* at 100nM drug concentration, we first calculated the MT tip diffusion coefficient and net-rates, as a measure of MT dynamic behavior, as a function of the fold changes in on- and off-rates of the dimers (dk_on_, dk_off_) (Fig. 11B and Fig. S2A in the supporting material). We found that ∼20- to 30-fold reduction of k_off_, as a result of stabilizing the lateral bond, and a ∼100 to 200 fold reduction of k_on_ at the poisoned tip, presumably as a result of conformational change of the longitudinal interface of the TC complex, recapitulates steady state zero net-rate and the stabilizing effects of colchicine at 100nM *in vitro* (Fig. 5D).

**Figure 11.**
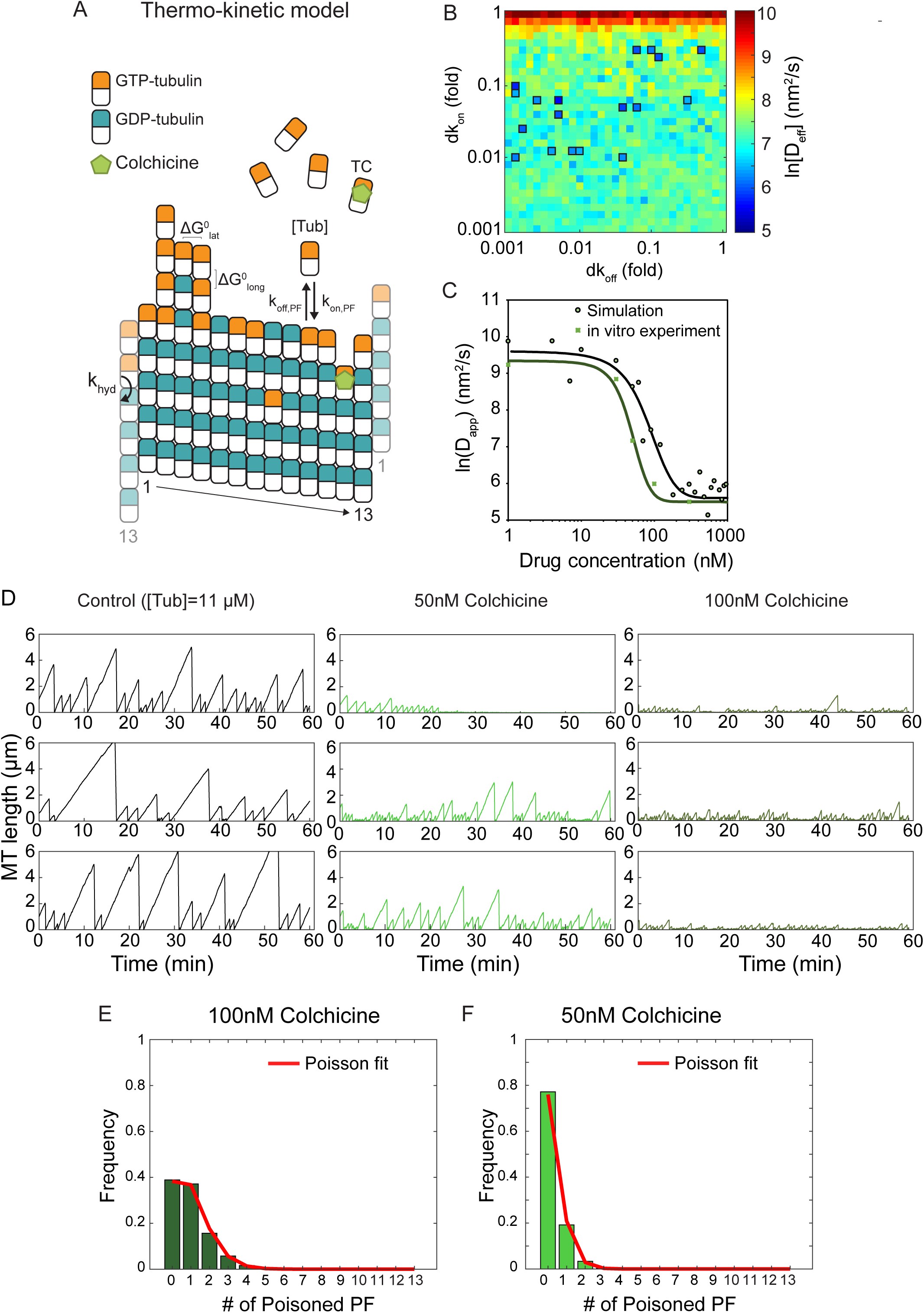
Poisson poisoning identified as the mechanism of kinetic stabilization by colchicine by thermo-kinetic modeling for microtubule self-assembly. (A) End-poisoning mechanism is shown in the thermo-kinetic model of MT assembly with model parameters. (B) Model-predicted MT plus-end diffusion coefficient (D_app_) as a function of lateral bond stabilization (k_off_ reduction) and end-poisoning (k_on_ reduction). All kinetic values were normalized to the base set for *in vitro* dynamic instability. The region with bold outlines indicates the parameter space consistent with experimentally observed stabilization with 100nM colchicine. *In vitro* parameter set were used in these simulations. (C) Model-predicted (circles) and *in vitro* experimental (square) MT assembly variance (D_app_) as a function of different colchicine concentrations. Here the drug kinetic parameters were chosen according to (A) and Fig. S2. (D) Example model-predicted MT life histories for control MTs and in the presence of 50nM and 100nM colchicine. MTs were each simulated for 60 minutes. (E) Poisson distribution of poisoned PFs at the tip of MTs, predicted by modeling MT assembly in the presence of 100nM and (F) 50nM colchicine.

We then examined the stabilization effect of colchicine by changing the fraction of TC complexes relative to free tubulin available (11 µM) at a specific free colchicine concentration (100nM) and the drug dissociation equilibrium constant (K_D_) from tubulin in the simulation (Fig. S2B in the supporting material). The diffusion coefficient results show that the model is mostly sensitive to the fraction of colchicine-bound tubulin rather than the affinity of the drug (K_D_). It is shown that at saturating and near-saturating conditions, where 80 to 100% of colchicine in solution is bound to free tubulin, the simulations would represent the *in vitro* poisoning conditions more closely. This further supports the hypothesis that the observed experimental MT stabilization effects were close to the saturation point after a 30-minute exposure to 100nM colchicine *in vitro*. Thus, all simulations were run assuming that all colchicine is fully bound to tubulin (100% saturation). We then chose the value of K_D_ based on the k_on_ and k_off_ of colchicine binding to tubulin as reported in previous literature (43, 58, 103) (42, 44) and our MT dynamics recovery experiments after colchicine washout *in vivo*. After determination of the drug parameter set (Table S1), we simulated several instances of an individual MT in the presence of TC complexes in the solution and examined MT dynamics as a function of colchicine concentrations (Fig. 11C). The MT diffusion values were in good agreement with *in vitro* experimental values. MT length vs. time histories (Fig. 8D) for control and colchicine treated MTs correlated well with the *in vitro* MT kymographs (Fig. S3 in the supporting material).

Thus, in contrast to previous MTAs, colchicine presented a unique MT poisoning mechanism with the fraction of poisoned PFs varying along different simulated MTs according to a Poisson distribution. As expected from *in vitro* experiments, colchicine at 100nM can block MT polymerization, with some nanomolar dynamics still going on. The number of poisoned PFs for this concentration for each time point in a 30-minute simulation time followed a Poisson distribution for several simulated MTs (Fig. 11E), with a mean value of 1 PF. Our simulation analysis indicated a minimum of 3 poisoned PFs (∼23% of total PFs) at the end of each simulation was required to create the kinetic stabilization effect, observed in experimental conditions. As for 50nM colchicine, the kinetic stabilization was weaker and only observed for shorter periods of time as predicted by *in vitro* experiments and the number of PFs poisoned varied between 1 and 3 (Fig. 11F). The Poisson poisoning effect at this concentration was occasionally repaired by addition of new dimers (Fig. S4 in the supporting material). Interestingly, at higher concentrations of colchicine (>100nM), although the number of poisoned PFs increases, events of further addition of tubulin and TC complexes were still observed and poisoned PFs were not fully blocked (∼100 fold reduction in k_on_). With higher fractions of TC complex in the PFs, the probability of repair and assembly recovery decreased, and a complete assembly inhibition was observed (Fig. S5 in the supporting material). This phenomenon is previously described as “copolymerization” of TC complexes in the MT lattice with tubulin (22, 23, 35).

Taken together, our thermokinetic modeling results, parametrized within the constraints of quantitative *in vivo* and *in vitro* experiments and molecular modeling, identifies a substoichiometric Poisson poisoning mechanism for colchicine effects on MT dynamics.

## Conclusions

In our study, we presented a multi-scale framework to examine the mechanism of action of colchicine on MT dynamics. Our results provided a connection between previously suggested mechanisms regarding colchicine binding to tubulin at different length-time scales as well as new insights into tubulin conformational changes following the drug binding. The significance of our analysis, in our view, is pinpointing a drug’s mechanism of action by connecting all the data from atomistic, molecular, and MT-level modeling to *in vitro* and *in vivo* experimental observations. Consequently, the identified mechanism of action is confined within several experimental and computational results and less dependent on the methodology of the experiments. Altogether, the results of our multi-scale analysis reveal that colchicine mainly binds to soluble form of tubulin due to the occlusion of the path to binging pocket in polymerized tubulin. After forming the TC complex, colchicine poisons the MT tips substoichiometrically, with the number of poisoned PFs varying along different MTs with a Poisson distribution. TC structure differs from a free tubulin significantly in the GTP-state, primarily in the conformations of the longitudinal and lateral bond-involved residues and intradimer bending angles. The conformational changes make the incorporation of additional dimers to a PF tip unfavorable (k_on_ reduction but not complete inhibition of association) and lateral bond more favorable (k_off_ reduction), which together kinetically stabilize MTs. In our view, our analysis offers compelling evidence for a copolymerization mechanism (22, 35) in which a TC complex has a lower affinity for tubulin polymerization rather than absolute blockage of assembly at the poisoned tip.

A number of potential limitations need to be considered in the current study. First, the molecular dynamics simulations investigated only the structure of colchicine bound to DARPin-decorated tubulins which by itself might affect the tubulin conformation compared to an undecorated pure tubulin structure. Despite this, we believe that our equilibration runs with water allowed the protein to relax its restraints due to DARPin binding. Second, our experiments did not measure an affinity for the drug colchicine directly, rather, relied on previous kinetic studies to estimate its binding affinity to tubulin. We note that our thermokinetic modeling recapitulated the stabilization effects of the drug, observed in experiments closely, with the estimated values of drug kinetics. Finally, we were unable to identify the underlying reason for lower efficacy of the drug in our cells compared to *in vitro* experiments. This was probably due to lower permeability of the drug by an active agent in the cell membrane which can be further investigated in future studies. However, we can still claim that the binding and unbinding of colchicine to tubulin is slow regardless of the lower permeability of the drug in our cells.

The findings of our study have two main implications for our understanding of MT assembly regulation by MTAs. First, as our understanding of the mechanism of action of colchicine has been enhanced, further clinical trials of the drug colchicine can be improved. In our opinion, clinically-acceptable low doses of colchicine can be effective in MT stabilization if the exposure time is adjusted based on the drug dose and the drug is administered locally. In contrast to other widely known MTAs such as paclitaxel and vinblastine that affect their target within minutes, exposure time to colchicine is a factor as important as the drug concentration when examining the poisoning effects. This further supports the results of a recent study on the successful blockage of migration and proliferation *in vitro* and *in vivo* with doses as low as 5nM and incubation up to 24 hours (18). In addition, colchicine can be combined with another MTA with different mechanism of action to tackle the problem of resistant cell lines. Second, our approach allows the replacement of an MTA or a self-assembled protein of interest, to be fully characterized for its effect on normal cell functions. Thus, all the colchicine-derivatives and CSIs, which have recently gained interest in treating multi-drug resistant cell lines (104–108), can be potentially tested for their efficacy and toxicity on MT dynamics and cell functions and compared to the narrow therapeutic window of colchicine.

## Author Contributions

Conceptualization, M.H. and D.J.O.; Methodology, M.H., and D.J.O.; Software, M.H. and D.E.; Validation, M.H., M.B.; Analysis, M.H., M.B., D.E. and D.J.O.; Writing – Original Draft, M.H. and D.J.O.; Writing – Review & Editing, all authors; Visualization, M.H., D.E.; Supervision, D.J.O.; and Funding Acquisition, D.J.O.

## Acknowledgements

The authors thank Dr. Brian Castle for his helpful discussions in preparing the paper. This study was supported by National Institutes of Health under award number R01-AG053951 and the Institute for Engineering in Medicine (IEM) award at the University of Minnesota to DJO. The authors acknowledge the Extreme Science and Engineering Discovery Environment (XSEDE), Bridges system at the Pittsburgh Supercomputing Center (PSC) through allocation MCB160060, and the Minnesota Supercomputing Institute (MSI) at the University of Minnesota for providing resources that contributed to the research results reported within this paper.

