## Supporting material for "Poisson poisoning as the mechanism of action of the microtubule-targeting agent colchicine"

### Supporting Tables

#### Table S1. Thermokinetic model *in vitro* parameter set

| Parameter | Definition | Value | Reference |
|---|---|---|---|
| $\Delta G_{lat}^0$ | Lateral bond free energy | -5 $k_B$T | Hemmat *et al.* (2019); Gardner *et al.* (2011) |
| $\Delta G_{long}^0$ | Longitudinal bond free energy | -7.2 $k_B$T | Castle *et al.* (2018) Gardner *et al.* (2011) |
| $\Delta\Delta G^0$ | Energetic penalty of GDP-tubulin | +3.5 $k_B$T | Hemmat *et al.* (2020) bioRxiv submission doi: 2020.01.07.897439 |
| [Tub] | Free tubulin concentration | 5.6 µM | Schek *et al.* (2007); Gardner *et al.* (2011) |
| $k_{on,PF}$ | On-rate constant | 6 $\mu M^{-1}s^{-1}PF^{-1}$ | Gardner *et al.* (2011) |
| $k_{hyd}$ | Hydrolysis rate constant | 0.2 $s^{-1}$ | Model constrained; Coombes *et al.* (2013) |
| $\sigma_1$, $\sigma_2$ | One- and two-neighbor on-rate penalty | 2, 10 | Castle *et al.* (2013) |
| $k_{on,\ colch.}$ | Colchicine on-rate constant to tubulin | 200 $M^{-1}s^{-1}$ | Garland *et al.* (1975) Banerjee *et al.* (1992) Diaz *et al.* (1991) |
| $k_{off,\ colch}$ | Colchicine off-rate from tubulin | $5 \times 10^{-6}$ $s^{-1}$ | Current study Garland *et al.* (1975) Diaz *et al.* (1991) |
| $KD_{colch}$ | Colchicine dissociation constant | 25 nM | Garland *et al.* (1975) Banerjee *et al.* (1992) Diaz *et al.* (1991) |

#### Table S2. Average length and radius of the identified channels connecting the surface and buried active site of GDP-tubulin.

| | Average Channel length (Å) | Average bottleneck radius (Å) | Colchicine entrance (%) |
|---|---|---|---|
| Tub (R) | 9.1 ± 0.8 | 3.7 ± 0.2 | 83 |
| TC (R) | 24 ± 3 | 3.2 ± 0.3 | 75 |
| $Tub_s$ (L) | 35 ± 2 | 1.4 ± 0.2 | 11 |

### Supporting Figures

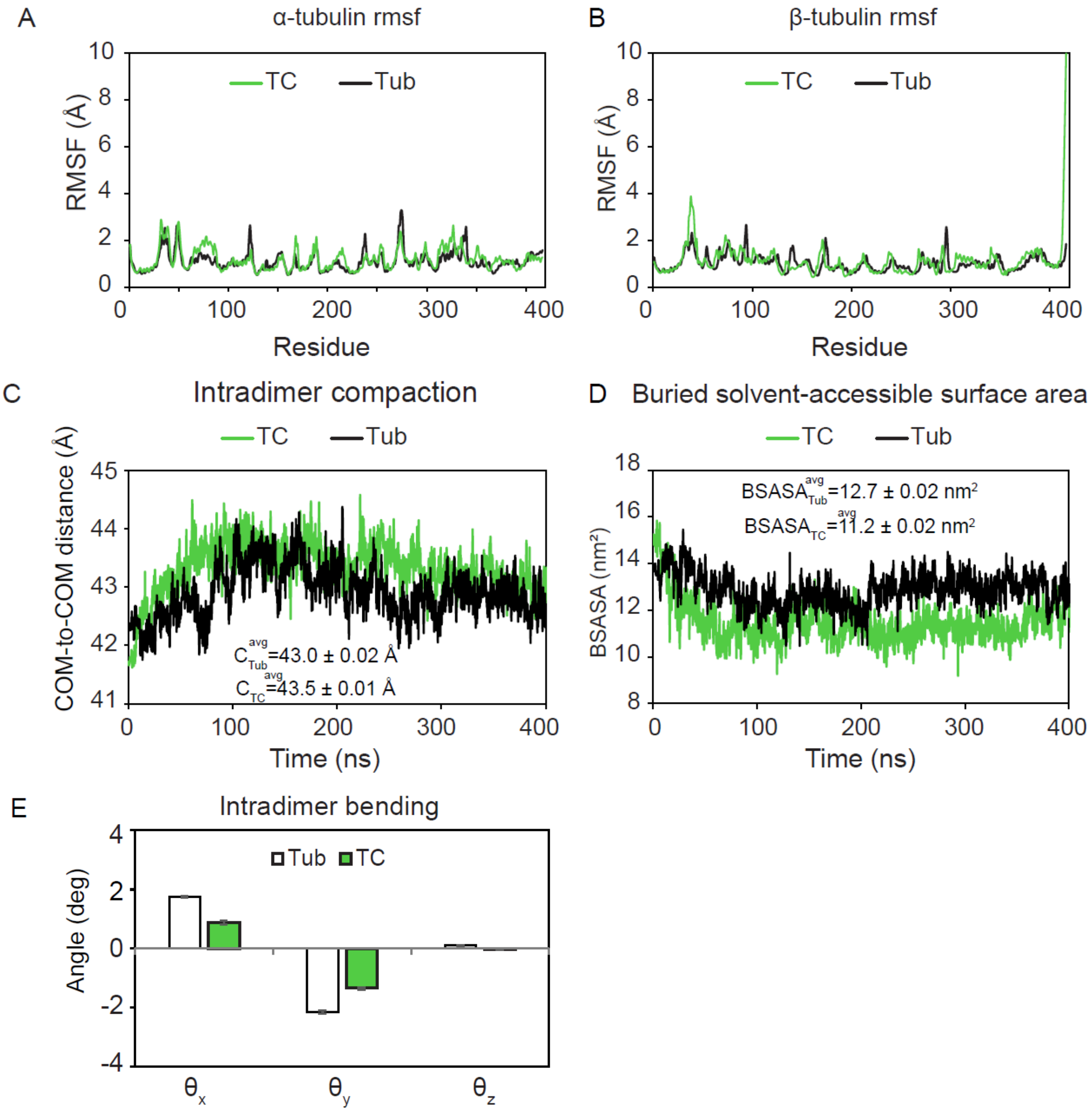


**Figure S1.** Molecular dynamics simulations showed conformational changes caused by colchicine binding in tubulin in GDP-state. (A) Average RMSF values of α- and (B) β-tubulin BD) for GDP TC and Tub complexes indicate dimer flexibility is not significantly changed upon drug binding. (C) Intradimer compaction, estimated from COM-to-COM distance of α- and β-subunits, is lower for GDP-TC complex compared to Tub. (D) Hydrophobic interactions, estimated from BSASA, are weaker in GDP-TC complex compared to Tub due to a larger intradimer spacing. (E) Intradimer bending angles are lower on average for TC complex compared to Tub due to limited flexibility in motion of the residues. Error bars are average of the boot-strapped data ± SEM .

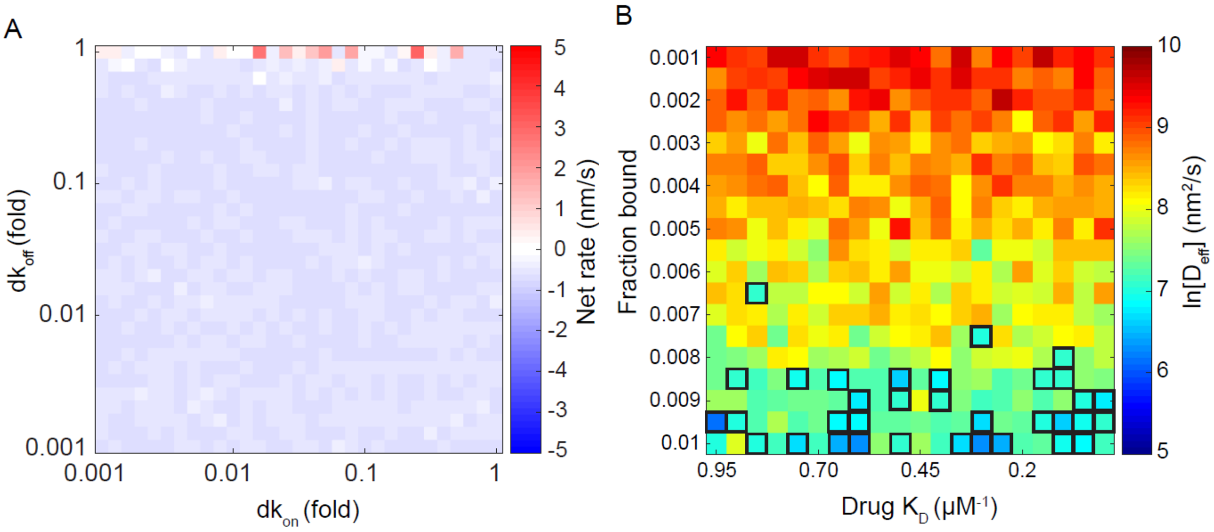


**Figure S2.** Model-predicted MT dynamics indicates the drug kinetic parameter set consistent with experimental results. (A) Net-rates as a function of fold changes in association ($k_{on}$) and dissociation ($k_{off}$) rates due to drug binding. Zero net-rates are almost observed for all parameter set used in the simulations. (B) MT tip apparent diffusion is predicted as a function of fraction of TC: tubulin in solution and drug affinity. Bold borders indicate consistent diffusion values with *in vitro* experiments for 100nM colchicine.

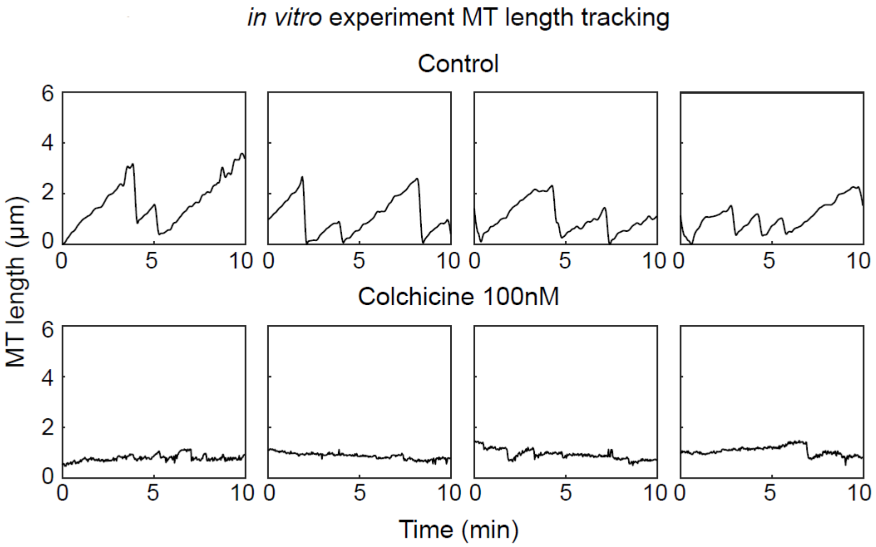


**Figure S3.** Examples of semiautomated tracking of MT dynamics *in vitro* experiments for control and 100nM colchicine-treated MTs.

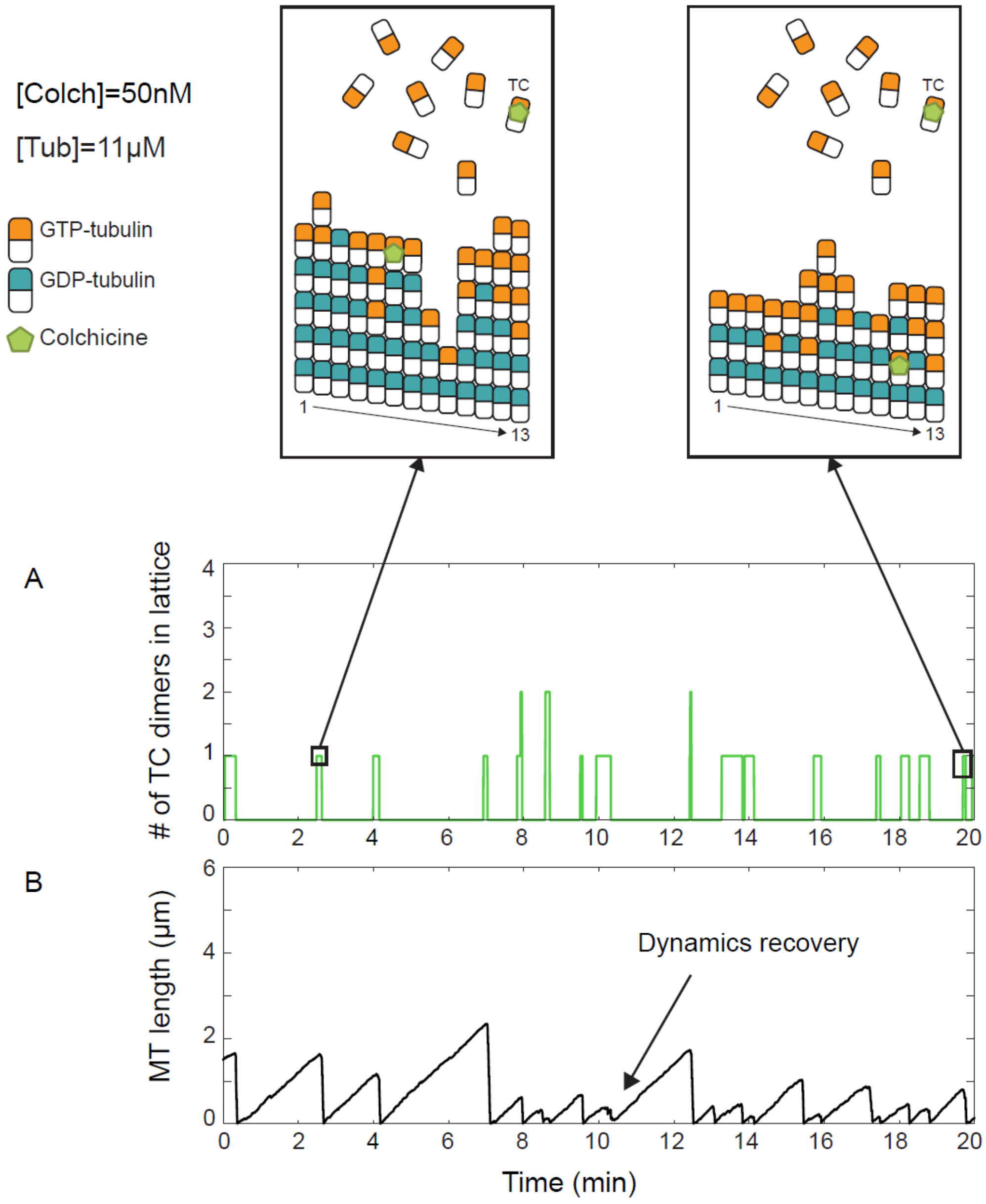


**Figure S4.** Poisoning mechanism for low colchicine concentration (50nM) predicted by our model shows occasional dynamic recovery by further addition of tubulin dimers. (A) Number of TC dimers incorporated into the lattice as a function of simulation time. Panels demonstrate examples of PF tips and TC distribution at an individual MT at time points near the beginning of simulation (left) and end of simulation (right). (B) MT length as a function of simulation time is shown. Arrowhead points to an occasional dynamic recovery when the TC complex was buried down the lattice by further dimer addition.

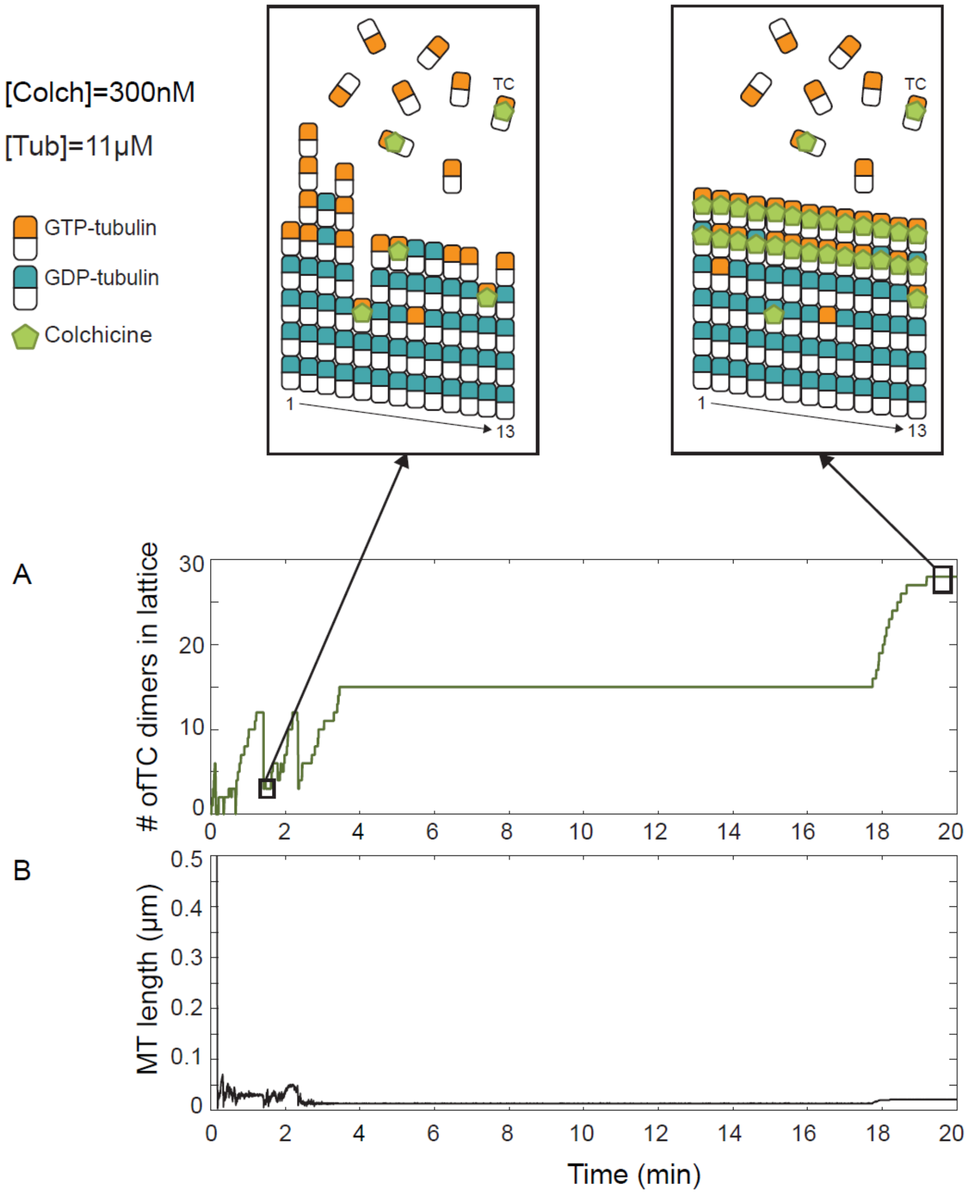


**Figure S5.** Poisoning mechanism for higher colchicine concentration (300nM) predicted by our model indicates assembly inhibition and copolymerization of a TC complex with itself and tubulin dimers. (A) Number of TC dimers incorporated into lattice as a function of simulation time. Panels demonstrate examples of PF tips and TC distribution at an individual MT at time points near the beginning of simulation (left) and end of simulation (right). (B) MT length as a function of simulation time is shown. Complete inhibition of MT assembly is observed after ~3 minutes of simulation.

**Supporting movies**

**Movie S1.** Microtubules in control LL-CPK1α cells show dynamic behavior. The time-lapse movie was recorded after 30 minutes incubation of the cells with fresh media. A total of 60 frames were collected at 1-s intervals.

**Movie S2.** Microtubules in 600nM colchicine-treated LL-CPK1α cells show kinetic stabilization. The time-lapse movie was recorded after 30 minutes exposure of the cells to the drug. A total of 60 frames were collected at 1-second intervals.

**Movie S3.** Control LL-CPK1α cells show normal migration and proliferation after media exchange. A total of 60 frames were collected at 10-minute intervals.

**Movie S4.** LL-CPK1α cells recovered after washout of 300nM colchicine show normal migration and proliferation. A total of 60 frames were collected at 10-minute intervals.

**Movie S5.** EB1 comets and their decay as a marker of hydrolysis in LLCPK1-EB1 cells. A total of 60 frames were collected at 1-second intervals.

**Movie S6.** EB1 comets and their decay as a marker of hydrolysis in LLCPK1-EB1 cells treated with 300nM colchicine for 24 hours. A total of 60 frames were collected at 1-second intervals.
